# Engineering of chimeric polyketide synthases using SYNZIP docking domains

**DOI:** 10.1101/489120

**Authors:** Maja Klaus, Alicia D. D’Souza, Aleksandra Nivina, Chaitan Khosla, Martin Grininger

## Abstract

Engineering of assembly line polyketide synthases (PKSs) to produce novel bioactive compounds has been a goal for over twenty years. The apparent modularity of PKSs has inspired many engineering attempts in which entire modules or single domains were exchanged. In recent years, it has become evident that certain domain-domain interactions are evolutionarily optimized, and if disrupted, cause a decrease of the overall turnover rate of the chimeric PKS. In this study, we compared different types of chimeric PKSs in order to define the least invasive interface and to expand the toolbox for PKS engineering. We generated bimodular chimeric PKSs in which entire modules were exchanged, while either retaining a covalent linker between heterologous modules or introducing a non-covalent docking domain- or SYNZIP domain-mediated interface. These chimeric systems exhibited non-native domain-domain interactions during intermodular polyketide chain translocation. They were compared to otherwise equivalent bimodular PKSs in which a non-covalent interface was introduced between the condensing and processing parts of a module, resulting in non-native domain interactions during the extender unit acylation and polyketide chain elongation steps of their catalytic cycles. We show that the natural PKS docking domains can be efficiently substituted with SYNZIP domains and that the newly introduced non-covalent interface between the condensing and processing parts of a module can be harnessed for PKS engineering. Additionally, we established SYNZIP domains as a new tool for engineering PKSs by efficiently bridging non-native interfaces without perturbing PKS activity.

## Introduction

Multimodular PKSs are large megasynthases that produce polyketides in an assembly line manner. The prototypical example is the 6-deoxyerythronolide B synthase (DEBS), which harbors six modules and produces the macrocyclic core of the antibiotic erythromycin (Fig. 1A). More than a thousand such assembly line PKSs have been discovered to date, each presumably synthesizing a distinct polyketide natural product.^1^ As many polyketide compounds exhibit strong biological activities, assembly line PKSs are interesting targets for engineering. Over the years many studies have attempted to either rationally engineer PKSs by domain exchanges,^2,3^ whole module exchanges,^4,5^ active site mutagenesis,^6–8^ or using evolutionary principles.^9,10^ Beyond traditional considerations of substrate specificity, a major constraint on engineering assembly line PKSs arises from the specificity of domain-domain interactions. Most notably, turnover of the assembly line requires that the acyl carrier protein (ACP) of each module must interact with all enzymatic domains within the module as well as the ketosynthase (KS) of the downstream module. For example, the ACP domain of Module 1 of DEBS interacts with two distinct KS domains during chain elongation and chain translocation (Fig. 1B).

**Figure 1.**
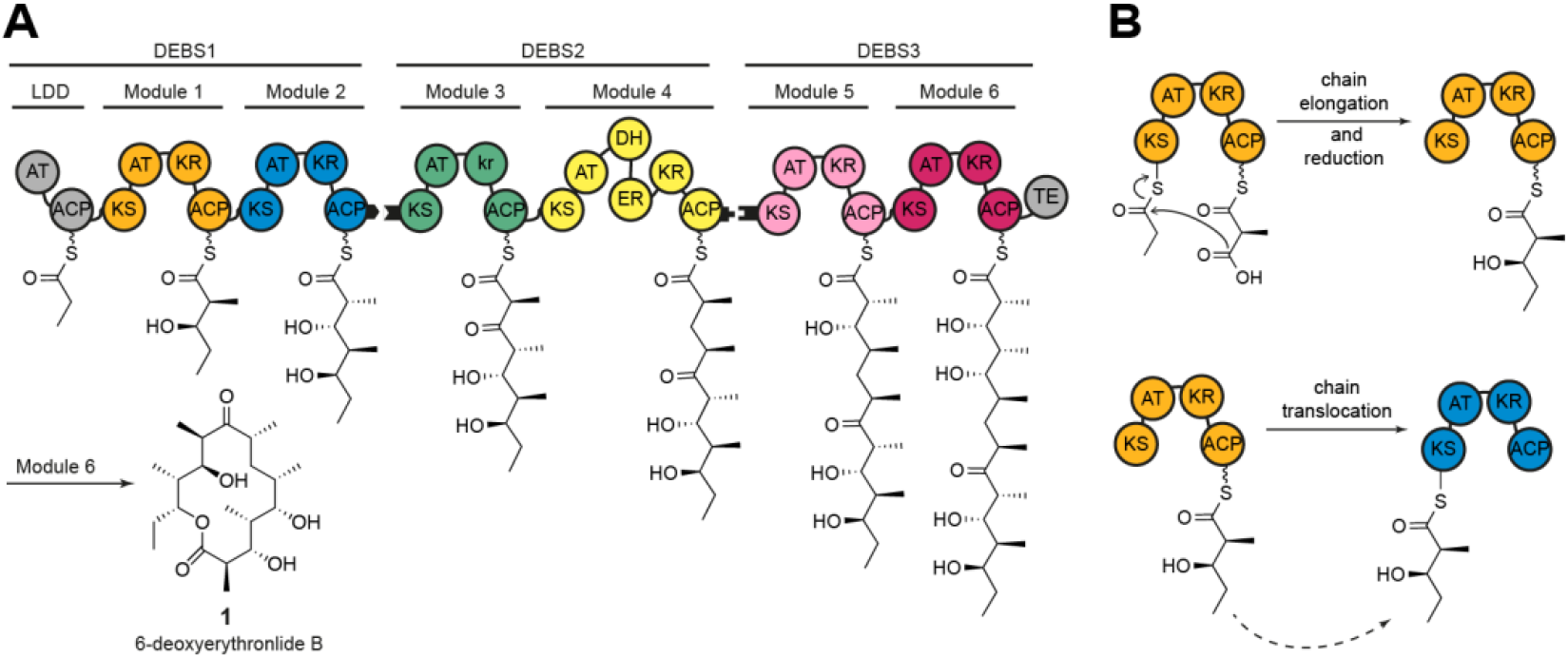
Schematic architecture of DEBS. A) The three polypeptides (DEBS1-3), the encoded modules (M1-M6), as well as the loading didomain (LDD), the thioesterase domain (TE), and the final product (**1**) are depicted. Polyketide intermediates are shown as attached to the respective acyl carrier protein (ACP). Black tabs depict docking domains. Domain annotations are as follows: AT – acyltransferase, KS – ketosynthase, ACP – acyl carrier protein, KR – ketoreductase, DH – dehydratase, ER – enoylreductase. Module 3 has a ketoreductase-like domain (denoted in lowercase) that lacks NADPH-dependent oxidoreductase activity but harbors C-2 epimerase activity. B) The intramodular chain elongation and intermodular chain translocation steps are shown for M1. The ACP of this module carries the α-carboxyacyl-chain (methylmalonyl-ACP) or the (2*S*,3*R*)-2-methyl-3-hydroxy-diketide condensation product.

Several studies have focused on the exchange of intact PKS modules to retain the inherent modularity of PKSs.^4,5,11^ To mediate communication between separate modules, PKS derived docking domains were installed at the N- and C-termini of heterologous modules. These docking domains are often relatively large and interact with weak affinities (K_D_ ≈ 20-100 μM).^12–14^ Chimeric PKSs harboring intact heterologous modules retain native domain-domain interactions within individual modules, but introduce a non-native ACP-KS interface in the chain translocation reaction (intermodular chain translocation interface),^15,16^ and therefore suffer from low product titers *in vivo*^4^ and decreased turnover rates *in vitro*^5^. Alternatively, structurally stable chimeric PKSs could also be engineered to preserve the chain translocation interface while modifying the intramodular chain elongation interface, by fusing the N-terminal condensing part (comprised of the KS and AT domains) of one module to the C-terminal processing part (harboring auxiliary enzyme(s) and ACP domain) of another. However, such possibilities may be constrained by KS-ACP specificity during polyketide chain elongation.^17,18^ There remains a need for a general method to access catalytically efficient chimeric assembly line PKSs.^19^

In this study, we explore the utility of heterospecific, high-affinity (K_D_ ≈ 10 nM^20,21^) coiled-coil interaction domains, termed SYNZIPs,^20,21^ for engineering chimeric PKS modules comprised of an N-terminal condensing part (KS-AT) and a C-terminal (DH-ER-KR)-ACP processing part derived from different modules. The non-covalent, SYNZIP domain-mediated interface was anticipated to be minimally invasive to a PKS module and as such could be harnessed for PKS engineering. This assumption was mainly built on structural data of PKSs and the related fatty acid synthases (FASs), where minimal contact is observed between the condensing and processing parts,^22–24^ thereby allowing extensive rotational movement of these two parts^25,26^. Furthermore, genetic analysis suggested an evolutionary relationship between the upstream (DH-ER-KR)-ACP parts and the downstream KS domains, indicating that introduction of an intramodular chimeric PKS interface between the AT domain and the downstream processing part may be beneficial as it preserves the evolutionary conserved intermodular interface.^27,28^ This in turn would enhance the modularity of the multienzyme system, allowing for numerous combinations in the generation of chimeric PKSs. Our findings suggest that a SYNZIP-mediated intramodular chimeric interface presents an effective alternative to recombining intact modules, especially in situations where the kinetic barrier of an engineered chain translocation step dominates that of chain elongation.

## Results and Discussion

### The use of SYNZIP domains for engineering chimeric PKS modules

As a test case for the utility of SYNZIP domains, we used a previously characterized bimodular PKS derived from DEBS (Fig. 1A) that is comprised of three proteins: LDD(4), (5)M1(2), (3) M2-TE.^29^ In this system, the numbers in parenthesis denote the origins of matching docking domains that are fused to the N- or C-termini of these three proteins. For example, LDD(4) refers to the loading didomain to which the docking domain from the C-terminus of DEBS Module 4 has been fused. Previous studies have shown that the AT-KR linker can be generally cleaved at a well-defined site without structural perturbations to either the N-terminal or C-terminal domains,^3,30^ but with a significant kinetic penalty to the chain elongation step^17^. Accordingly, DEBS M1 was genetically cleaved at this site (Fig. 2A). The choice of SYNZIP domains to introduce at this junction was guided by the length and orientation of the resulting coiled-coil. Among the best characterized orthogonal SYNZIP pairs are SYNZIP1+SYNZIP2 and SYNZIP3+SYNZIP4 (hereafter annotated as SZ1+SZ2 and SZ3+SZ4) both interacting in a parallel orientation. While both pairs form highly stable heterodimers (k_off, sz1+sz2_ ≈ 7.8 x 10^-4^ s^-1^, k_off, SZ3+SZ4_ ≈ 1.4 x 10^-2^ s^-1^),^31^ we chose the SZ3+SZ4 pair, as the SZ1+SZ2 coiled-coil is predicted to be longer.^21^ Throughout this study, SZ3 was fused to the C-terminus of one target protein, while SZ4 was fused to the N-terminus of another target protein. Being approximately 60 Å long, the SZ3+SZ4 coiled coil is predicted to be of the same length as the entire KS dimer (Fig. 2B). Because we used a (GGSG)_2_-linker to connect the SYNZIP domains to the PKS module fragments, we assumed the conformational flexibility of the ACP would be sufficient to allow for interaction with the KS and AT domain of the reconstituted module.

**Figure 2.**
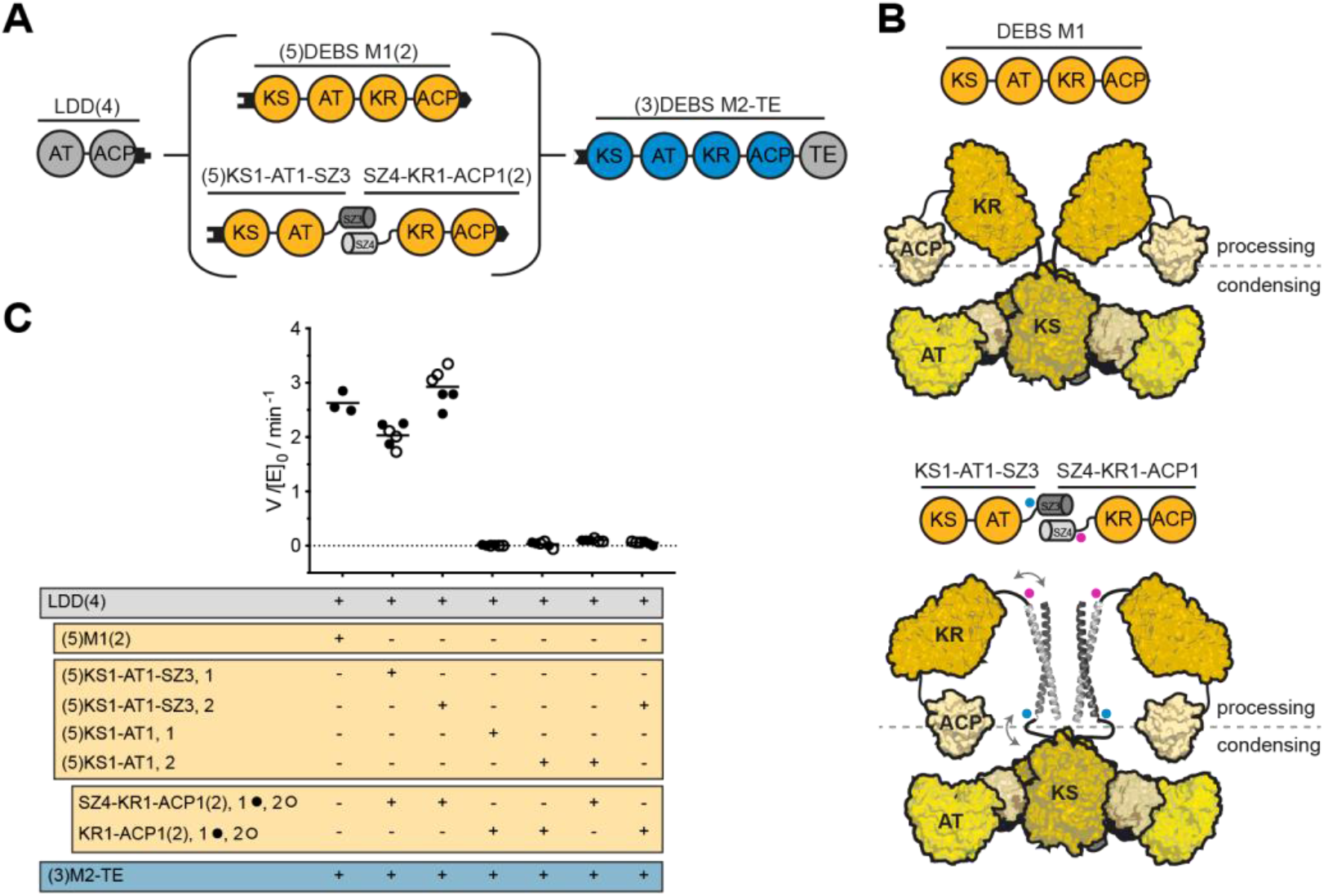
Use of SYNZIP domains in the design of catalytically efficient PKSs harboring a split module. A) Design of a bimodular DEBS derivative comprised of LDD(4), intact (5)M1(2) or a split version thereof, and (3)M2-TE. B) Model of DEBS M1 with the natural AT-KR linker (top) and the SYNZIP-containing variant (bottom). This model was built based on SAXS analysis of DEBS M3^32^. Fusion sites of the SYNZIP domains to either the AT or KR are indicated by blue/pink dots, illustrating the parallel orientation of the heterospecific coiled-coil. An eight-residue flexible Gly-Ser linker is used to connect the SYNZIP domain to the PKS protein (see protein sequence Table S2). C) Turnover rates of bimodular PKSs employing M1 or a split M1. Except for intact M1, at least two independently purified protein preparations were evaluated (1 or 2): each preparation of the N-terminal part is shown in a separate column, whereas preparations of the C-terminal part are indicated as black and white dots. All initial rate data was obtained at 2 μM enzyme concentration and non-limiting concentrations of propionyl-CoA, methylmalonyl-CoA, and NADPH. Measurements were performed in triplicates and the grand mean is indicated.

Using a UV_340_ spectrophotometric assay, we measured the turnover rate of the bimodular PKS in the presence of either intact M1, split M1 bridged by SYNZIP domains, split M1 without SYNZIP domains, or split M1 in which only one of the SYNZIP domains needed to form the heterodimer was present (Fig. 2C). Remarkably, the turnover rates of the PKSs harboring intact M1 or SYNZIP-bridged M1 did not differ significantly (Fig. 2C, first three columns), while omission of either or both SYNZIP domains led to a predicted large drop in turnover rates (Fig. 2C, last four columns). These results indicate that a SYNZIP-bridged module can achieve comparable catalytic activity to an intact one. By introducing a non-covalent SYNZIP pair at the junction site of the condensing and processing part of a module, it is thus possible to achieve a high effective molarity of the α-carboxyacyl-ACP substrate for the KS-catalyzed chain elongation step.

As the cleavage at the AT-KR junction site most deleteriously affects the KS:ACP interaction during chain elongation,^17,30^ we next sought to compare the rates of chain elongation between an intact M1 and a split M1 with or without SYNZIP domains. For this purpose, we used a radioisotope labeling assay in which [1-^14^C]-propionyl-CoA is used to prime the LDD, thereby allowing measurement of the occupancy of the N-terminal and C-terminal fragments of Module 1 with radiolabeled polyketide intermediates. First, we performed a set of assays in the absence of methylmalonyl-CoA, the co-substrate required for chain elongation. As shown in Fig. 3A, the N-terminal (KS-AT) fragment of M1 was labeled rapidly (within 1 min) regardless of the presence or absence of a SYNZIP domain. Presumably this labeling was due to efficient translocation of the propionyl moiety from LDD to the KS active site. As expected, in the absence of SYNZIP domains, no labeling of the ACP-containing C-terminal protein was observed, nor did the labeling intensity of the intact M1 protein exceed an average occupancy of one equivalent^33^ at any time. However, in the presence of SYNZIP domains, labeling of the C-terminal protein increased steadily over 10 min. The absence of methylmalonyl-CoA prevents chain elongation and transfer of the labeled growing polyketide chain to the C-terminal protein, which suggests that this labeling is indicative of transacylation of the propionyl moiety from the KS to the downstream ACP. This may arise from the greater flexibility of the ACP domain in the SYNZIP system compared to intact M1, thereby allowing access to the KS domain in an orientation that is precluded in a native module. This transacylation reaction might be more pronounced in the absence of the co-substrate for chain elongation.

While chain translocation from LDD to the KS domain of a split module was unaffected by the presence of a SYNZIP interface (Fig. 3A), upon the addition of methylmalonyl-CoA, chain elongation showed strong dependence on the SYNZIP domains (Fig. 3B). The ACP-containing C-terminal protein labeled rapidly (within one 1 min) and the combined labeling intensity of the two SYNZIP-fused proteins was similar to that observed for intact M1, suggesting that the effective molarity of the α-carboxyacyl-ACP species was comparable in the two cases. A turnstile model has recently been proposed to account for the ability of an unoccupied KS active site to discriminate between the occupied versus unoccupied states of its partner ACP domain.^33^ Assuming that module M1 saturates at an average occupancy of one radiolabeled acyl chain per module, our data argues that the turnstile mechanism has been preserved in SYNZIP-bridged Module 1, notwithstanding significant differences in module geometry and flexibility in the two cases (Fig. 2B).

**Figure 3.**
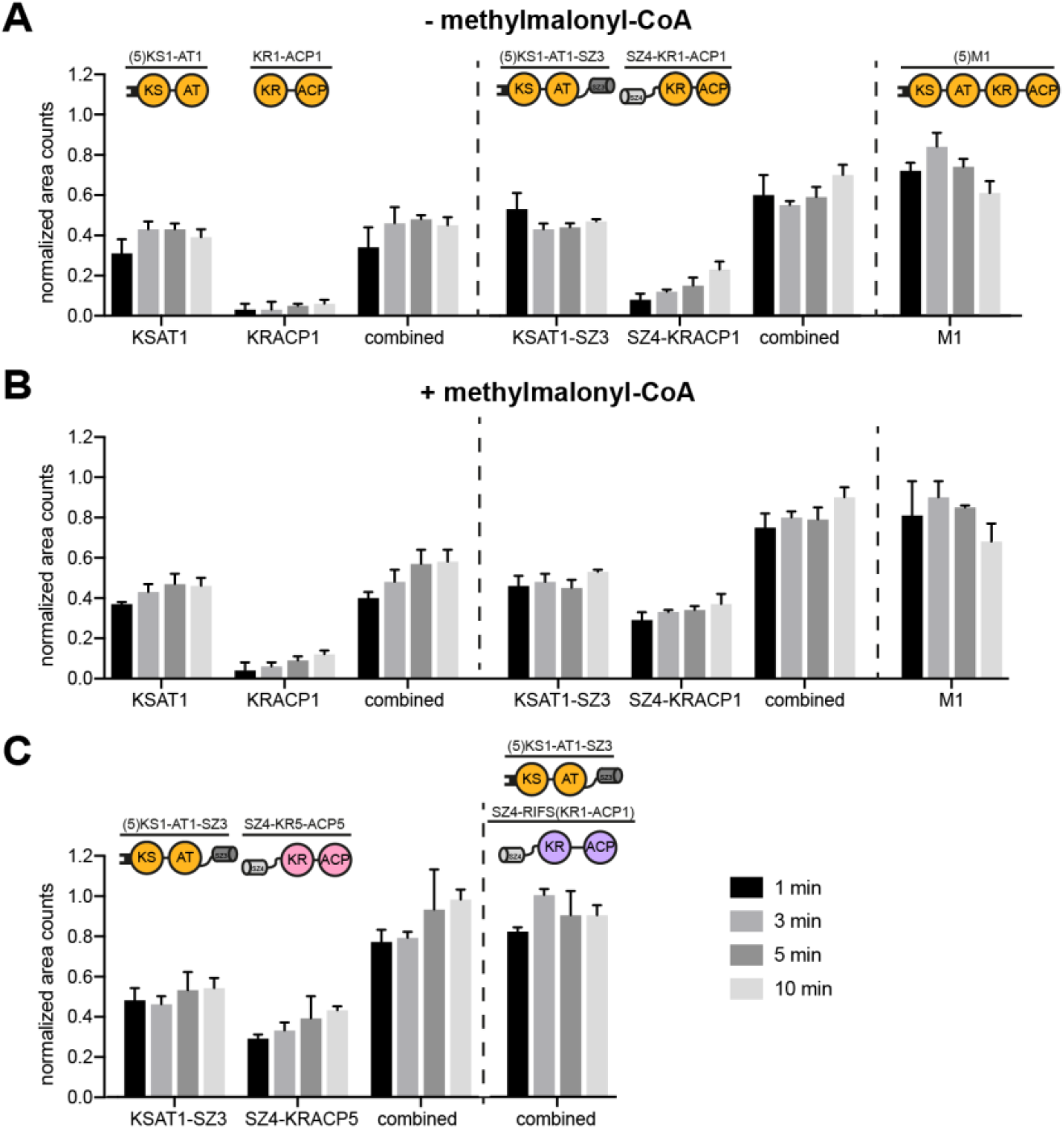
Comparison of chain translocation and elongation rates of intact versus split modules. The occupancy of individual proteins by growing polyketide chain precursors was measured in the absence (A) or presence (B) of methylmalonyl-CoA, and in modules containing chimeric KS:ACP interfaces (C, with methylmalonyl-CoA). All measurements included LDD(4) plus the designated split/intact module combinations. Labeling of the different proteins was quantified at different time points (1, 3, 5, and 10 min) and the resulting counts were normalized based on the maximal occupancy. Combined counts of both the KS and ACP containing proteins are also presented for split-module systems. In the case of RIFS Module 1, the N- and C-terminal fragments could not be separated by SDS-PAGE gel; hence only combined counts are reported (panel C). All measurements were performed in triplicates using 2 μM enzyme concentration and non-limiting concentrations of propionyl-CoA, methylmalonyl-CoA, and NADPH.

Encouraged by the above results, we sought to examine whether SYNZIP domains could facilitate labeling between the KS-AT fragment of Module 1 and the ACP domain of a noncognate module, indicative of efficient chain elongation across a chimeric interface. Specifically, we measured the occupancy of the SZ4-fused C-terminal (KR-ACP) fragments of DEBS Module 5 and rifamycin synthase (RIFS) Module 1 (Fig. 3C). In both cases the combined counts of the N- and C-terminal proteins were comparable to those of the reference intact module. In light of the observed ability of a KS-bound polyketide chain to transacylate to the ACP in a split-module system harboring SYNZIP domains (Fig. 3A), it was not possible to conclude from this data alone whether chain transacylation or chain elongation kinetics were being enhanced by the SYNZIP interface in these chimeric modules. However, given that the apparent condensation rate of the reference system (Fig. 3B) was much faster than the transacylation rate (Fig. 3A), the strong labeling of the chimeric C-terminal proteins was assumed to originate mostly from elongation rather than transacylation.

Taken together, the above data highlighted the potential utility of SYNZIP domains as tools to engineer non-covalently recombined chimeric modules at the AT:KR interface.

### Design and activity of chimeric PKSs using different domain-domain interfaces

Following this initial assessment, we extended our evaluation of SYNZIP domain-mediated engineering to a larger set of bimodular chimeric PKSs, in which the non-cognate domain interface corresponds to the intermodular chain translocation step of the PKS catalytic cycle (Fig. 1B). This set of PKSs had previously been studied with intact heterologous modules interfaced by non-covalent docking domains.^4,5,11^ We constructed three types of bimodular PKSs: using non-covalent docking domains (Fig. 4A), a SYNZIP-interfaced system (Fig. 4B), and a combination of covalent fusions and SYNZIP domains (Fig. 4C). Earlier analysis of a subset of analogous covalently fused chimeric PKSs had revealed quantitatively distinct properties compared to non-covalently fused systems.^15^ As detailed below, careful examination of our findings can reconcile these quantitative differences.

As reported previously,^5^ when non-covalent docking domains were engineered at the intermodular interface between DEBS M1 and various downstream modules, only the wild-type bimodular PKS with DEBS M2 as the acceptor achieved high turnover rates. All bimodular chimeric PKSs exhibited significantly decreased turnover rates (Fig. 4A). When SYNZIP domains were installed at the intermodular interface, the cognate DEBS M1-M2 system achieved similar turnover rates as the respective docking domain-interfaced one (Fig. 4B). In addition, the turnover rate of the system harboring DEBS M6 as an acceptor increased significantly in the presence of SYNZIP domains, while the turnover rates of chimeric PKSs harboring DEBS M3 or DEBS M5 did not change appreciably (Fig. 4B). In the third type of bimodular PKSs, the split Module 1 is bridged by SYNZIP domains and the downstream acceptor module is covalently fused to the C-terminal fragment of Module 1 (Fig. 4C). Similarly to the second design, the cognate DEBS M1-M2 system was unaffected. However, chimeric systems harboring DEBS M3 and M6 showed increased turnover rates compared to the analogous docking domain-interfaced systems (Fig. 4C).

**Figure 4.**
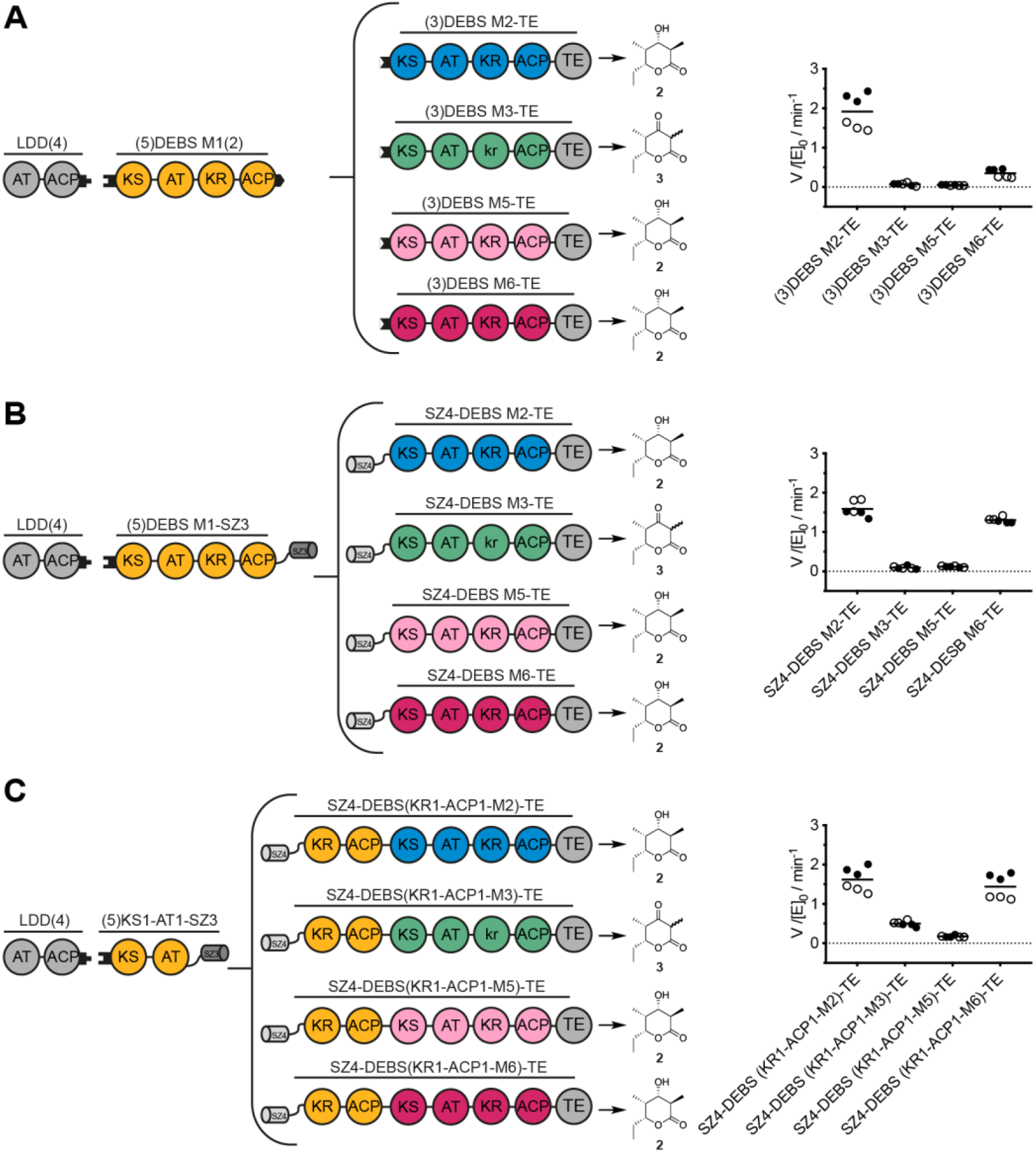
Turnover rates of bimodular chimeric PKSs harboring a chimeric chain translocation interface. Design and turnover rates of bimodular chimeric PKSs. LDD(4) was always used as a separate protein, whereas DEBS M1 and variable acceptor modules were interfaced using either non-covalent docking domains (A), SYNZIP domains (B) or a combination of covalent fusions and SYNZIP domains (C). Construct design and the predicted triketide lactone products are shown next to the measured turnover rate of each system. All initial rate data was obtained at 2 μM enzyme concentration and non-limiting concentrations of propionyl-CoA, methylmalonyl-CoA, and NADPH. In all cases two independently purified protein preparations were evaluated (data shown as black and white dots). Each set of measurements was performed in triplicates and the grand mean is indicated. LC-MS analysis was performed after overnight incubation to verify product identity (Fig. S4).

Data on bimodular systems (see Fig. 2 and Fig. 4) suggests that SYNZIP-interfaced systems can enhance the turnover rates of some chimeric PKSs. SYNZIP interfaces appear to be comparable to covalent linkages in this respect, with the added benefit of mitigating the need to express and purify large proteins such as intact bimodules.^29^ Because bridging the split Module 1 with SYNZIP domains is minimally deleterious to protein structure (see Fig. 4C), and because chimeric modules showed elongation kinetics approaching that of the wild-type system (see Fig. 3C), a new and superior strategy for engineering chimeric PKSs may be to install non-cognate interfaces within modules while in turn preserving the cognate module-module interfaces.

In order to test this hypothesis, the PKS design outlined in Fig. 5 was utilized. The N-terminal fragment is identical to the one used in the previous assays and is bridged to the C-terminal fragment by SYNZIP domains (Fig. 4C). However, the location of the chimeric interface in the C-terminal fragment has been shifted from elongation to translocation: the processing parts of DEBS M5 and RIFS M1 are fused to their cognate downstream modules DEBS M6 and RIFS M2, respectively. Kinetic analysis of both chimeras revealed significantly attenuated turnover rates (Fig. 5). LC-MS analysis after overnight incubation revealed the correct masses for the expected products of the reference system and the PKS with DEBS modules 5 and 6, yet not with RIFS modules 1 and 2 (Fig. S4). The precise mechanism for these functional impairments was not established, but attenuated rates may be caused by impaired substrate recognition imposed by binding pocket or domain-domain interface specificity.^6,34^ Alternatively, intramodular transacylation of the polyketide intermediate from the KS to the ACP could also present a barrier for polyketide chain growth, as illustrated for Module 1 (Fig. 3B); this could be especially deleterious for chimeric modules with inherently low turnover rates. This side reaction might be suppressed by redesigning the SYNZIP system to limit the conformational variability of ACP, for example by shortening the coiled coil, rigidifying Gly-Ser linkers, or using lower-affinity docking domains.

**Figure 5.**
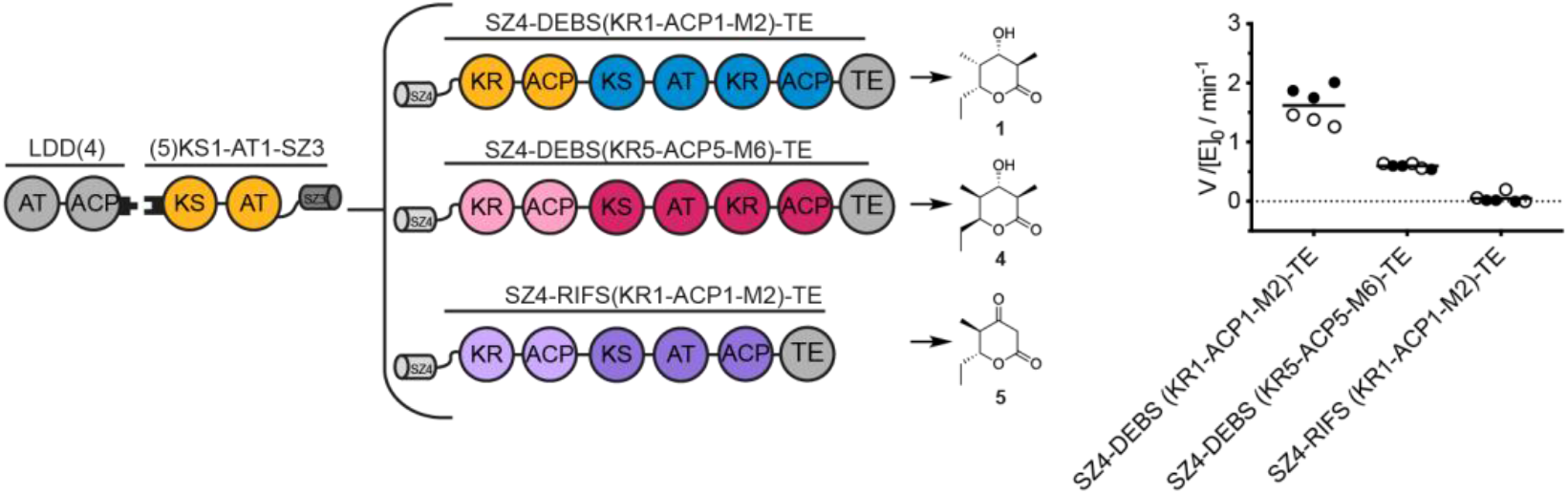
Turnover rates of bimodular chimeric PKSs harboring a chimeric chain elongation interface. Bimodular PKSs comprised of LDD(4), (5)KS1-AT1-SZ3 and variable SZ4-KR_n_-ACP_n_-Module_n+1_-TE constructs. Construct design and the predicted triketide lactone products are shown next to the measured turnover rate of each system. All initial rate data was obtained at 2 μM enzyme concentration and non-limiting concentrations of propionyl-CoA, methylmalonyl-CoA, malonyl-CoA (RIFS M2 is malonyl specific), and NADPH. Turnover analysis was performed on two individually purified proteins per construct (back and white dots). Measurements were performed in triplicates and the grand mean is indicated.

## Conclusion

Several different chimeric PKS designs involving SYNZIP domains installed at either a non-native chain translocation or chain elongation interface were systematically compared in this study. Whereas the chain translocation interface is well established, the benefit of docking domains at the chain elongation interface had not been investigated prior to this study. Our findings suggest that SYNZIP domains have considerable potential, especially in instances where proximity generated by a high-affinity linkers can improve ACP:KS interactions. In contrast, other chimeric PKSs cannot be optimized by trading one non-native protein-protein interface for another in a straightforward manner.^35^ We conclude that both interfaces are equivalently suited for PKS engineering, but that the resulting chimeric PKSs are often kinetically impaired, requiring further optimization. Overall, our findings showcase the utility of SYNZIP domains as a new tool in PKS engineering that can replace traditionally used PKS docking domains while also mitigating the need for expressing and purifying excessively large PKS proteins. Their small size, high affinity interaction, and the multitude of existing orthogonal pairs makes them ideal candidates to bring non-natively interacting PKS modules/domains into close proximity. As they alter the geometry of a PKS module, they could also be used to study orientation, flexibility and conformational restraints of domains.

## Experimental Procedures

### Reagents

CloneAmp HiFi PCR Premix was from Takara. Restriction enzymes were from New England Biolabs. All primers were synthesized by Elim Biopharm or Sigma Aldrich. For DNA purification, the GeneJET Plasmid Miniprep Kit and the GeneJET Gel Extraction Kit from Thermo Scientific were used. Stellar Competent Cells were from Takara and One Shot^®^ BL21 (DE3) Cells were from Invitrogen. All chemicals for buffer preparations were from SigmaAldrich. Isopropyl-β-D-1-thiogalactopyranoside (IPTG), kanamycin sulfate, and carbenicillin were from Gold Biotechnology. LB-Miller Broth for cell cultures was from Fisher Scientific. Ni-NTA affinity resin was from MC Lab. For anion exchange chromatography the HiTrapQ column was from GE Healthcare. For all SDS-PAGE analyses SDS-PAGE Mini Protean TGX precast gels were from Bio-Rad (4-20% and 7.5%). For protein concentration Amicon Ultra centrifugal filters were from Millipore. Coenzyme A (CoA), reduced β-nicotinamide adenine dinucleotide 2’-phosphate (NADPH), sodium propionate, propionic acid, methylmalonic acid, malonic acid, and magenesium chloride hexahydrate were from SigmaAldrich. Adenosine-5’-triphosphate (ATP) was from Teknova. Reducing agent tris(2-carboxyethyl)phosphine (TCEP) was from Thermo Scientific. UVette cuvettes (2 mm x10 mm path) were from Eppendorf.

### Plasmids

Plasmids harboring genes encoding individual PKS modules were either generated in this study via Infusion Cloning, site directed mutagenesis, or reported previously. Table S1 specifies the primer sequences and the resulting plasmids. Plasmids sequences were verified by DNA sequencing (Elim Biopharm or Sigma Aldrich). Protein sequences are presented in Table S2.

### Bacterial cell culture and protein purification

All PKS proteins were expressed and purified using similar protocols. For *holo*-proteins (where the ACP domain is post-translationally modified with a phosphopantetheine arm) *E. coli* BAP1 cells^36^ were used as the host. All proteins contained a C-terminal His_6_-tag for purification. Cultures were grown on a 1 L scale at 37 °C to an OD_600_ of 0.3, whereupon the temperature was adjusted to 15 °C. At OD_600_ of 0.6 protein production was induced with 0.1 mM IPTG, and the cells were grown for another 18 h. Cells were harvested by centrifugation at 5000 *g* for 8 min and lysed by sonication in a buffer consisting of 50 mM sodium phosphate, 10 mM imidazole, 450 mM NaCl, and 10% glycerol, pH 7.6. The cell debris was removed by centrifugation at 25,000 g for 1 h. The supernatant was incubated on Ni-NTA agarose resin (2 mL resin per liter of culture) at 4 °C for 1 h. Afterwards, the mixture was applied to a Kimble-Kontes Flex column and washed with the above lysis buffer (20 column volumes). Additional washing was performed with 10 column volumes wash buffer (50 mM phosphate, 25 mM imidazole, 300 mM NaCl, 10 % glycerol, pH 7.6). Proteins were eluted with 6 column volumes 50 mM phosphate, 500 mM 10% glycerol, pH 7.6. Using a HitrapQ column, additional purification was performed by anion exchange chromatography on an ÄKTA FPLC system. Buffer A consisted of 50 mM phosphate, 10% glycerol, pH 7.6, whereas buffer B contained 50 mM phosphate, 500 mM NaCl, 10% glycerol, pH 7.6. The yields of all protein per liter of culture are noted in Table S3. Enzymes MatB, SCME and PrpE were purified as decribed.^29,33^ Protein concentrations were determined with the BCA Protein Assay Kit (Thermo Scientific). Samples were stored as aliquots at –80 °C until further use.

### SEC analysis

To determine the purity of the purified protein after ion exchange chromatography, samples of each protein were analyzed on an ÄKTA FPLC system using a Superose 6 Increase 10/300 GL column with a buffer containing 50 mM phosphate, 500 mM NaCl, 10% glycerol, pH 7.6. For protein purity by SDS-PAGE and SEC profiles refer to Fig. S1-S3.

### PKS Enzymatic Assays

PKS enzymatic assays were performed according to published procedures.^5^

### Liquid Chromatography-Mass Spectrometry Analysis of Polyketides

Dried samples were reconstituted in 100 μL methanol, separated on a ZORBAX Extend C18 column (Agilent, 1.8 um, 2.1 × 50 mm), connected to an Agilent Infinity 1290 II HPLC over a 6 min linear gradient of acetonitrile from 5% to 95% in water, and subsequently injected into a 6545 QTOF mass spectrometer. Reduced and unreduced triketide products were located by searching for the theoretical *m/z* for the [M+Na]^+^- and [M+H]^+^-ion. Unreduced triketides: [M+Na]^+^ = 193.080 and [M+H]^+^ = 171.098, reduced triketides [M+Na]^+^ = 195.100 and [M+H]^+^ =173.118.

### ^14^C-Radioisotopic SDS-PAGE Labeling Assay

[1-^14^C]-Propionate was used to interrogate intramodular chain elongation within intact DEBS M1, broken variants thereof, and chain elongation between chimeric KS-AT and KR-ACP proteins. A 10x substrate mix was generated by mixing 0.4 mM [1-^14^C]-propionate, 2 mM methylmalonate, 2.4 mM CoA, 3.5 μM PrpE, 2 μM MatB, and 4 μM SCME in a reaction containing 400 mM sodium phosphate, pH 7.2, 5 mM TCEP, 10 mM MgCl_2_, and 8 mM ATP and incubated for 1 h. To quantify the efficiency of channeling of the labeled propionyl group successively from the LDD to either intact M1 or broken variants thereof, the substrate mixture was diluted ten-fold into individual reaction mixtures containing 2 μM LDD(4) and 2 μM of either intact M1 of 2 μM of each a KS1-AT1 construct and a KR-ACP construct in the presence of 400 mM sodium phosphate, pH 7.2, 5 mM TCEP, and 0.75 mM NADPH. By minimizing the amount of labeled propionyl-CoA in the reaction mixture, non-specific transfer of the radiolabel directly from the LDD to an acceptor module was minimized. At specified time-points, reactions were quenched by addition of Laemmli buffer, and samples were separated via 7.5% SDS-PAGE at 200 V for 44 min. The gel was washed with water for 5 min and stained with SimplyBlue™ SafeStain (Invitrogen) for 20 min. Following destaining with water for 5 min, the gel was mounted on a filter paper and dried *in vacuo* for 2 h using a Bio-Rad 543 Gel Dryer. The dried gel was imaged for 20 min lane by lane to quantify ^14^C on individual protein bands using a Rita Star TLC Analyzer (Raytest). Peaks were integrated to quantify the radiolabel bound to each protein.

## Supporting information

## Acknowledgments

We thank Mislav Oreb for the plasmid carrying the SYNZIP domains, Kevin Erazo for help with the protein purification, and the CheM-H Metabolic Chemistry Analysis Center for LC-MS analysis.

