## Supplementary material for "Engineering of chimeric polyketide synthases using SYNZIP docking domains"

**Supporting Information contents:**

Figure S1: Analysis of protein purity by SDS-PAGE and SEC – single module acceptors
Figure S2: Analysis of protein purity by SDS-PAGE and SEC – covalent fusion acceptors
Figure S3: Analysis of protein purity by SDS-PAGE and SEC – DEBS M1 and its derivatives
Figure S4: LC-MS analysis of products produced by bimodular chimeric PKSs newly generated in this study.
Table S1: Plasmids and primers used in this study
Table S2: Amino acid sequences of proteins used in this study
Table S3: Yields of proteins used in this study

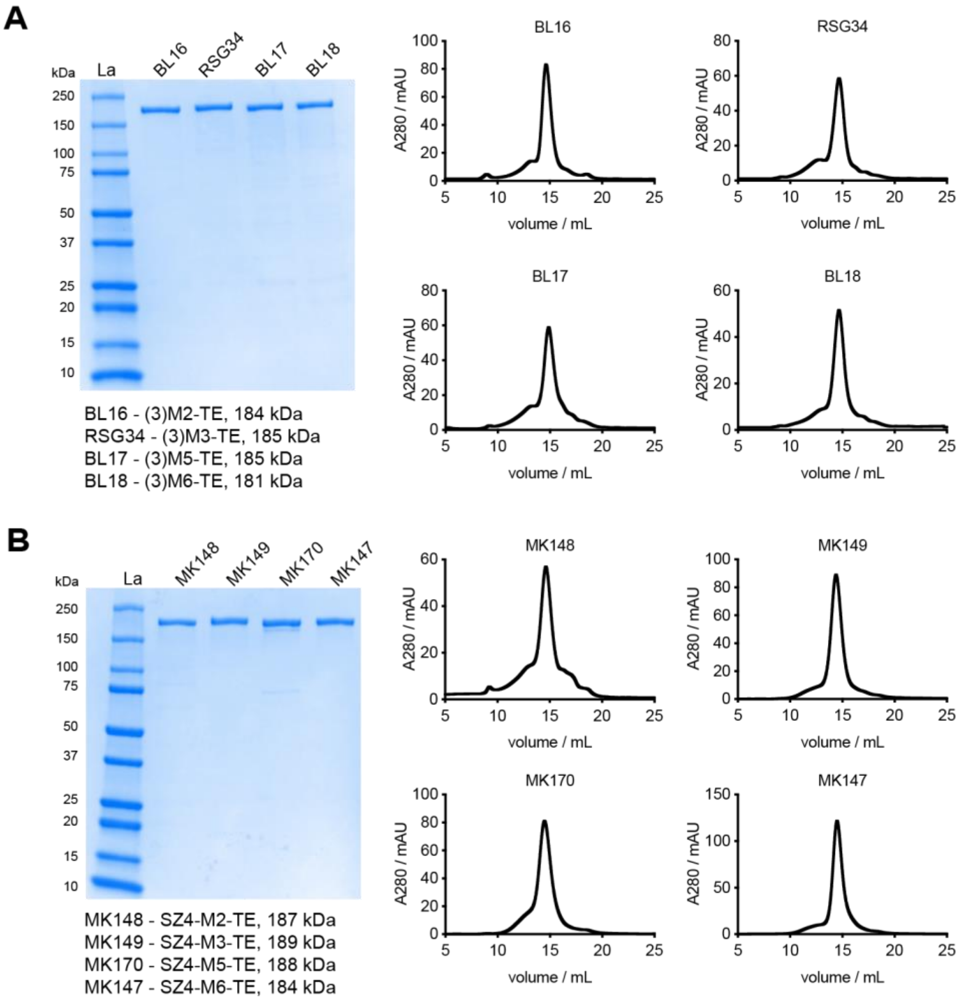


**Figure S1. Analysis of protein purity by SDS-PAGE and SEC – single module acceptors**. Acceptor proteins harboring a DEBS docking domain (A), and in comparison, a SZ domain (B). Protein abbreviations and their molecular weights (MW) are indicated. All proteins are pure and eluted in a predominately single peak from SEC.

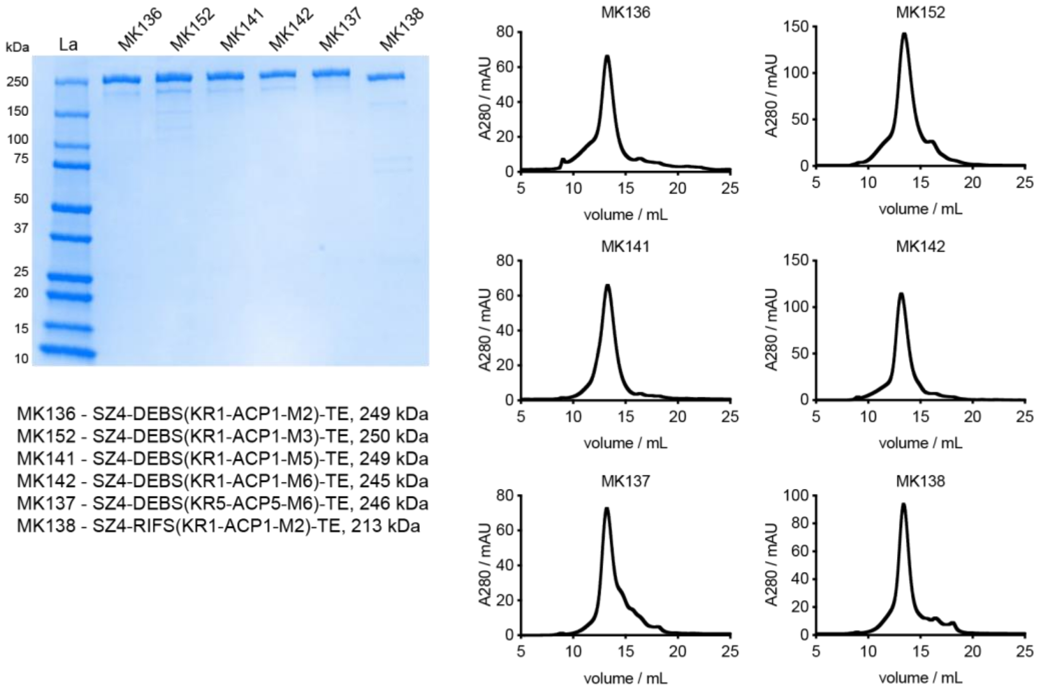


**Figure S2. Analysis of protein purity by SDS-PAGE and SEC – covalent fusion acceptors**. Acceptor proteins in which a KR-ACP fragment was fused to the acceptor module. Protein abbreviations and their MW are indicated. All proteins are pure and eluted in a predominately single peak from SEC.

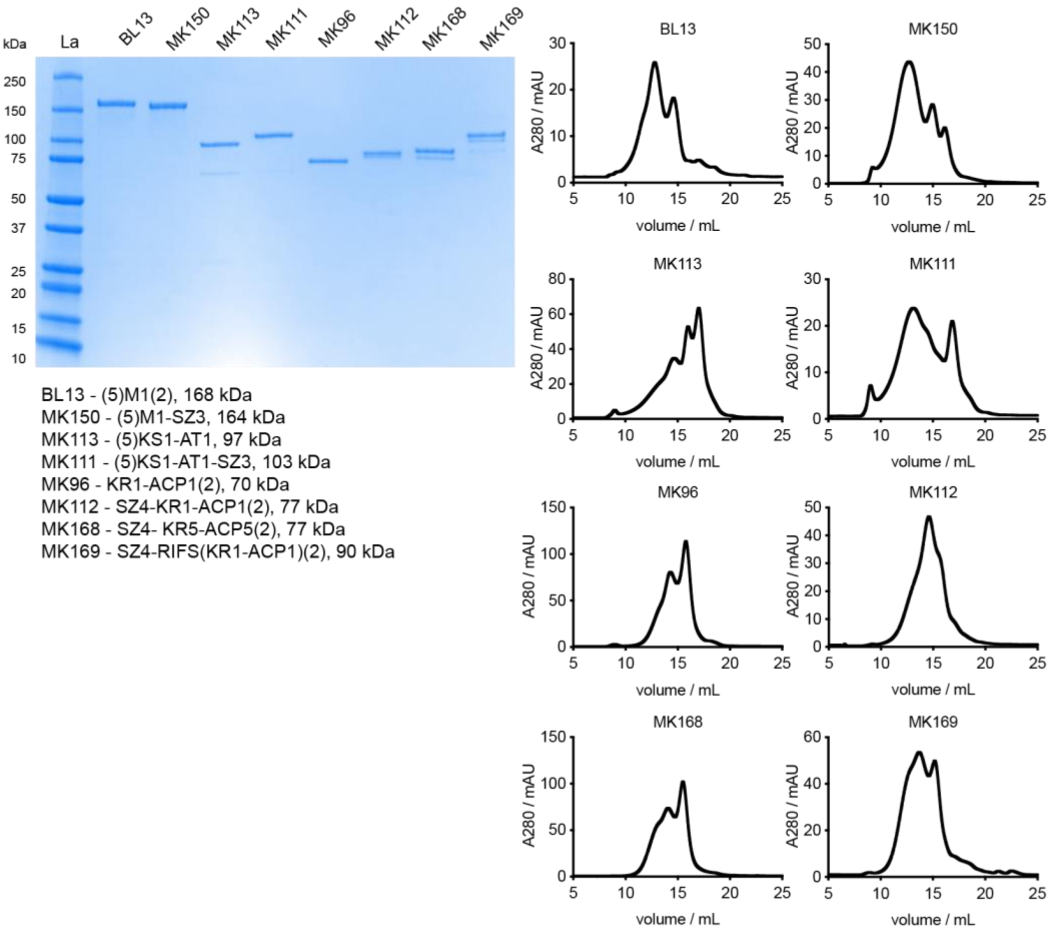


**Figure S3. Analysis of protein purity by SDS-PAGE and SEC – DEBS M1 and its derivatives**. DEBS M1 and its derivatives/fragments used in this study. If no indication is given the PKS domain/modules are derived from DEBS. Protein abbreviations and their MW are indicated. All proteins are pure, yet they appear as multiple oligomeric species based on SEC.

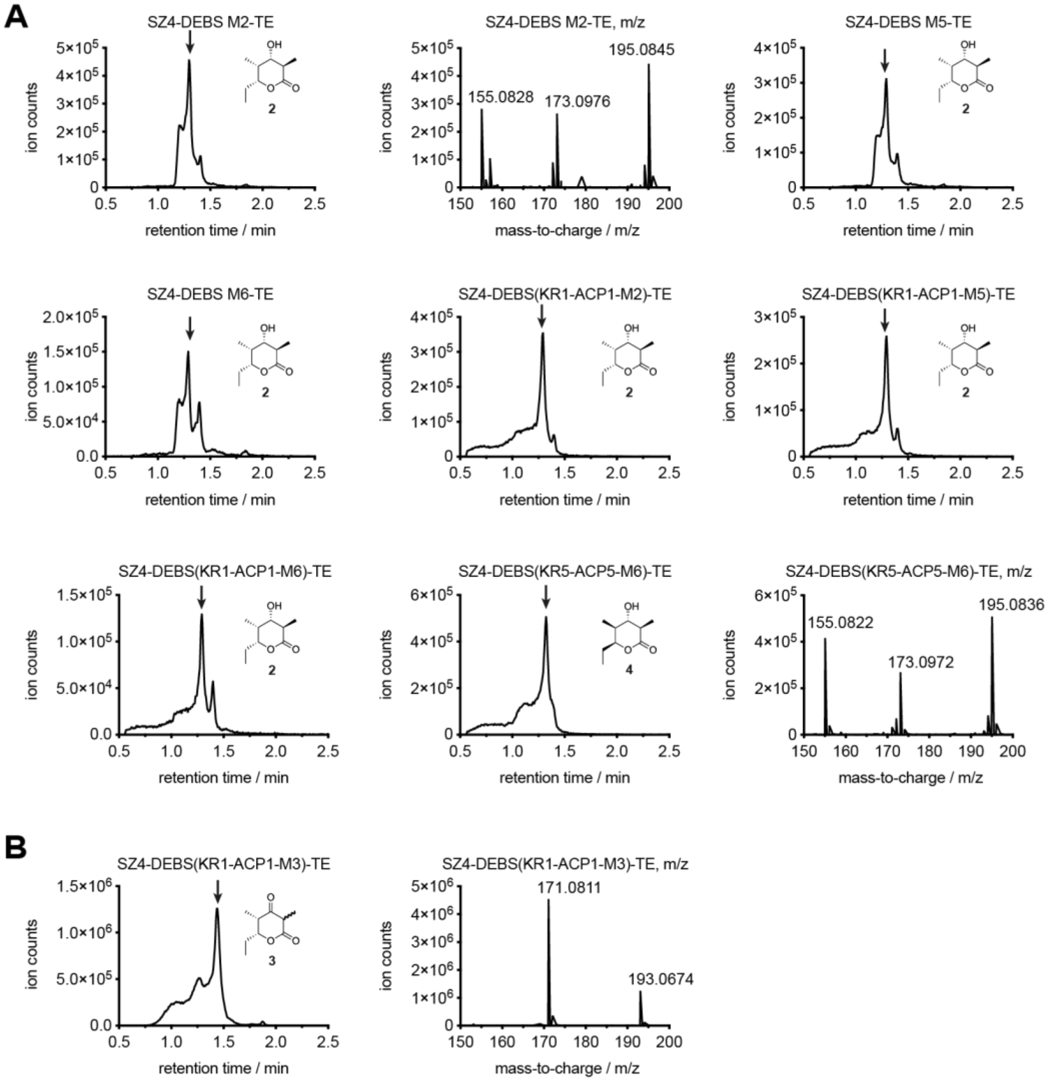


**Figure S4. LC-MS analysis of products produced by bimodular chimeric PKSs newly generated in this study.** A) Reduced triketide lactone products **2** and **4** ($C_9H_{16}O_3$, calculated MW 172.110) were detected in reaction mixtures containing SZ4-M2-TE, SZ4-M5-TE, SZ4-M6-TE, SZ4-KR1-ACP1-M2-TE, SZ4-KR1-ACP1-M5-TE, SZ4-KR1-ACP1-M5-TE, and SZ4-KR5-ACP5-M6-TE. B) Ketolactone **3** ($C_9H_{14}O_3$, calculated MW 170.090) was detected in reaction mixtures containing SZ4-KR1-ACP1-M3-TE. All PKS domain/modules are derived from DEBS. For all systems the extracted ion chromatograms were obtained by extraction of the $[M+Na]^+$ species and one chromatogram per compound is shown as an example. Labeled peaks from left to right correspond to $[M+H-H2O]^+$, $[M+H]^+$, and [M+Na]+ ions. The peak of interest is marked with an arrow based on its mass spectrum.

**Table S1: Plasmids and primers used in this study**. The cloning strategy is indicated for each construct. Some plasmids were only generated to function as intermediate constructs of the given cloning strategy and were not used for protein purification. If no indication is given the PKS domain/modules are derived from DEBS.

| Plasmid Encoded Protein | Cloning Method | Cloning Fragments | Primer Name | Primer Sequence 5'-3' | Template |
|---|---|---|---|---|---|
| pMK80 - ACP1-M5-TE, used to generate pADD02 | Infusion | 80_V1 | P-MK254 | CGCCTGGCTGGGCGGAGGG | pBL17 |
| | | | P-MK239 | CATATGTATATCTCCTTCTTAAAGTTAAACAAAATTA | |
| | | 80_V2 | P-MK250 | GAGCCGATCGCGATCGTCGGCATGGCGT | pBL17 |
| | | | P-MK255 | CACGCCCTGCGCGTACGCCTCGCC | |
| | | 80_V3 | P-MK256 | GCGTACGCGCAGGGCGTG | pBL17 |
| | | | P-MK253 | CCCTCCGCCCAGCCAGGCG | |
| | | 80_I | P-MK236 | AAGAAGGAGATATACATATGCTGGCGTCGCTGCCCG | pCK7 |
| | | | P-MK242 | CCGCGGGTGGGCGCGCTCGAGCACCACCACCACCACCACTGAGATC | |
| pMK81 - ACP1-M6-TE, used to generate pMK93 | Infusion | 81_V1 | P-MK257 | GACCCGATCGCGATCGTCGGCATGGC | pBL18 |
| | | | P-MK258 | CACGCCGTGCACGTGCGCCCGG | |
| | | 81_V2 | P-MK249 | GCGCACGTGCACGGCGTG | pBL18 |
| | | | P-MK239 | CATATGTATATCTCCTTCTTAAAGTTAAACAAAATTA | |
| | | 81_I | P-MK236 | AAGAAGGAGATATACATATGCTGGCGTCGCTGCCCG | pCK7 |
| | | | P-MK246 | CCGACGATCGCGATCGGGTCGACGGGGGCCGTGGTC | |
| pMK83 - ACP1-M2-TE, used to generate pADD03 | Infusion | 83_V1 | P-MK250 | GAGCCGATCGCGATCGTCGGCATGGCGT | pBL17 |
| | | | P-MK251 | GACACCGCGCGTGTGCGCGTCGG | |
| | | 83_V2 | P-MK252 | CGCACACGCGCGGTGTC | pBL16 |
| | | | P-MK253 | CCCTCCGCCCAGCCAGGCG | |
| | | 83_V3 | P-MK254 | CGCCTGGCTGGGCGGAGGG | pBL16 |
| | | | P-MK239 | CATATGTATATCTCCTTCTTAAAGTTAAACAAAATTA | |
| | | 80_I | P-MK236 | AAGAAGGAGATATACATATGCTGGCGTCGCTGCCCG | pCK7 |
| | | | P-MK242 | CCGCGGGTGGGCGCGCTCGAGCACCACCACCACCACCACTGAGATC | |
| pMK91- (5)KS1-AT1, used as a template for other cloning strategies | Infusion | 91_V | P-MK239 | CATATGTATATCTCCTTCTTAAAGTTAAACAAAATTA | pBL13 |
| | | | P-MK262 | TTTCCGCGCTGCGCTACCGCCTCGAGCACCACCACCAC | |
| | | 91_I1 | P-MK228 | GTGCAGCCCGTGATGTTCGCGGTCATGG | pBL13 |
| | | | P-MK261 | GTGGTGGTGGTGCTCGAGGCGGTAGCGCAGCGCGGAAACCTCGT | |
| | | 91_I2 | P-MK229 | CCATGACCGCGAACATCACGGGCTGCAC | pBL13 |

| | | | | | |
|---|---|---|---|---|---|
| | | | P-MK260 | GAAGGAGATATACATATGAGCGGTGACAACGGCATGACCGAGGAAAAG | |
| pMK93 - KR1-ACP1-M6-TE, missing an Asp in ACP1-KS2 linker | Infusion | 93_V | digestion of pMK81 with NdeI | | |
| | | 93_I | P-MK276 | AAGGAGATATACATATGGACGAGGTTTCCGCGCTG | pAYC59 |
| | | | P-MK277 | AGCGACGCCAGCATATGCGCGCCCACCCGCGGTTC | |
| pMK96 - KR1-ACP1(2) | Infusion | 96_V | P-MK239 | CATATGTATATCTCCTTCTTAAAGTTAAACAAAATTA | pBL13 |
| | | | P-MK62 | GGCACCGAGGTCCGGG | |
| | | 96_I | P-MK276 | AAGGAGATATACATATGGACGAGGTTTCCGCGCTG | pBL13 |
| | | | P-MK280 | CCGGACCTCGGTGCCGAGTTCGGCGGCCAGGT | |
| pMK111 - (5)KS1-AT1-SZ3 | Infusion | 111_V | P-MK309 | TCCGCCACCGGATCCGCCGAGCCAGACGCGCTCGC | pMK91 |
| | | | P-MK304 | CTGGCACACAAAAAGCTCGAGCACCACCACCAC | |
| | | 111_I | P-MK308 | GGATCCGGTGGCGGATCCGGTAACGAAGTTACAACACTTGAGAATGAC | pfRSZ |
| | | | P-MK303 | CTTTTTGTGTGCCAGTCTATTTCTCAAT | |
| pMK112 - SZ4-KR1-ACP1(2) | Infusion | 112_V | P-MK239 | CATATGTATATCTCCTTCTTAAAGTTAAACAAAATTA | pMK96 |
| | | | P-MK311 | TCCGGTGGCGGATCCGGTGACGAGGTTTCCGCGCTG | |
| | | 112_I | P-MK305 | GGAGATATACATATGCAGAAAGTGGCTGAATTGAAAAACAGA | pfRSZ |
| | | | P-MK310 | GGATCCGCCACCGGATCCGCCTTCAGCAACATCGTTCTCCAATCTG | |
| pMK113 - (5)KS1-AT1 (shorter than pMK91) | Infusion | 113_V | P-MK239 | CATATGTATATCTCCTTCTTAAAGTTAAACAAAATTA | pMK91 |
| | | | P-MK313 | GAGCGCGTCTGGCTCCTCGAGCACCACCACCAC | |
| | | 113_I | P-MK260 | GAAGGAGATATACATATGAGCGGTGACAACGGCATGACCGAGGAAAAG | pMK91 |
| | | | P-MK312 | GAGCCAGACGCGCTCGC | |
| pADD02 - KR1-ACP1-M5-TE, missing an Asp in ACP1-KS2 linker | Infusion | ADD02_V | digestion of pMK80 with NdeI | | |
| | | ADD02_I | P-MK276 | AAGGAGATATACATATGGACGAGGTTTCCGCGCTG | pAYC59 |
| | | | P-MK277 | AGCGACGCCAGCATATGCGCGCCCACCCGCGGTTC | |
| pADD03 - KR1-ACP1-M2-TE - missing an Asp in ACP1-KS2 linker | Infusion | ADD03_V | digestion of pMK83 with NdeI | | |
| | | ADD03_I | P-MK276 | AAGGAGATATACATATGGACGAGGTTTCCGCGCTG | pAYC59 |
| | | | P-MK277 | AGCGACGCCAGCATATGCGCGCCCACCCGCGGTTC | |
| pMK133 - KR1-ACP1-M2- | Quickchange | 133_QC | P-MK347 | GCGACCACGGCCCCCGTCGATGAGCCGATCGCGATCGTC | pADD03 |
| | | | P-MK348 | GACGATCGCGATCGGCTCATCGACGGGGGCCGTGGTCGC | |

| | | | | | |
|---|---|---|---|---|---|
| TE, complete ACP1-KS2 linker | | | | | |
| pMK134 - KR1-ACP1-M5-TE, complete ACP1-KS2 linker | Quickchange | 134_QC | P-MK347 | GCGACCACGGCCCCCGTCGATGAGCCGATCGCGATCGTC | pADD02 |
| | | | P-MK348 | GACGATCGCGATCGGCTCATCGACGGGGGCCGTGGTCGC | |
| pMK135 - KR1-ACP1-M6-TE, complete ACP1-KS2 linker | Quickchange | 135_QC | P-MK349 | GCGACCACGGCCCCCGTCGATGACCCGATCGCGATCGTC | pMK93 |
| | | | P-MK350 | GACGATCGCGATCGGGTCATCGACGGGGGCCGTGGTCGC | |
| pMK136 - SZ4-KR1-ACP1-M2-TE | Infusion | 136_V | digestion of pMK133 with NdeI | | |
| | | 136_I | P-MK351 | GAAGGAGATATACATATGCAGAAAGTGGCTGAATTGAAAAAC | pMK112 |
| | | | P-MK345 | AGCGACGCCAGCATATGCGCGCCCACCCGC | |
| pMK137 - SZ4-KR5-ACP5-M6)-TE | Infusion | 137_V | P-MK247 | GACCCGATCGCGATCGTC | pMK93 |
| | | | P-MK239 | CATATGTATATCTCCTTCTTAAAGTTAAACAAAATTA | |
| | | 137_I1 | P-MK351 | GAAGGAGATATACATATGCAGAAAGTGGCTGAATTGAAAAAC | pMK112 |
| | | | P-MK352 | GCCGGTGGGGATGGGACCGGATCCGCCACCG | |
| | | 137_I2 | P-MK353 | CCCATCCCCACCGGCG | pBL130 |
| | | | P-MK354 | GATCGCGATCGGGTCGTCGGCATCCTTCGGCAC | |
| pMK138 - SZ4-RIFS(KR1-ACP1-M2)-TE | Infusion | 138_V | P-MK358 | GAGATCGGCACCGCCGCGGCCGAGGAGCCGATCGCGATCGTC | pAJ21 |
| | | | P-MK239 | CATATGTATATCTCCTTCTTAAAGTTAAACAAAATTA | |
| | | 138_I1 | P-MK351 | GAAGGAGATATACATATGCAGAAAGTGGCTGAATTGAAAAAC | pMK112 |
| | | | P-MK355 | GGGCTCGGCGGGCTCACCGGATCCGCCACCG | |
| | | 138_I2 | P-MK356 | GAGCCCGCCGAGCCC | pAJ20 |
| | | | P-MK357 | GGCGGTGCCGATCTCGGCCGGCCGGTTCGCCGCCGATCCGAAGAGCTT GGCGCGCAG | |
| pMK141 - SZ4-KR1-ACP1-M5-TE | Infusion | 141_V | digestion of pMK134 with NdeI | | |
| | | 136_I | P-MK351 | GAAGGAGATATACATATGCAGAAAGTGGCTGAATTGAAAAAC | pMK112 |
| | | | P-MK345 | AGCGACGCCAGCATATGCGCGCCCACCCGC | |
| pMK142 - SZ4-KR1-ACP1-M6-TE | Infusion | 142_V | digestion of pMK135 with NdeI | | |
| | | 136_I | P-MK351 | GAAGGAGATATACATATGCAGAAAGTGGCTGAATTGAAAAAC | pMK112 |
| | | | P-MK345 | AGCGACGCCAGCATATGCGCGCCCACCCGC | |
| pMK146 - SZ4-MCS-H6, intermediate | Infusion | 146_all | P-MK386 | TGCGGCCGCAAGCTTACCGGATCCGCCACCGG | pMK112 |
| | | | P-MK387 | AAGCTTGCGGCCGCACTC | |

| | | | | | |
|---|---|---|---|---|---|
| vector for various cloning strategies | | | | | |
| pMK147 - SZ4-M6-TE | Infusion | 147_V | digestion of pMK146 with HindIII | | |
| | | 147_I | P-MK388 | CGGATCCGGTAAGCTTGACCCGATCGCGATCGTC | pMK137 |
| | | | P-MK389 | GTGCGGCCGCAAGCTTCGAATTCCCTCCGCCCAG | |
| pMK148 - SZ4-M2-TE | Infusion | 148_V | digestion of pMK146 with HindIII | | |
| | | 148_I | P-MK390 | CGGATCCGGTAAGCTTGAGCCGATCGCGATCGTC | pMK136 |
| | | | P-MK389 | GTGCGGCCGCAAGCTTCGAATTCCCTCCGCCCAG | |
| pMK149 - SZ4-M3-TE | Infusion | 149_V | digestion of pMK146 with HindIII | | |
| | | 149_I | P-MK391 | CGGATCCGGTAAGCTTGACCCGATCGCCATCGTC | pRSG34 |
| | | | P-MK389 | GTGCGGCCGCAAGCTTCGAATTCCCTCCGCCCAG | |
| pMK150 - (5)M1-SZ3 | Infusion | 150_V | P-MK394 | ACGACCGCGACCGGTTC | pMK111 |
| | | | P-MK389 | GTGCGGCCGCAAGCTTCGAATTCCCTCCGCCCAG | |
| | | 150_I | P-MK392 | GAACCGGTCGCGGTCGT | pBL13 |
| | | | P-MK393 | GCCGAGTTCGGCGGCC | |
| pMK152 - SZ4-KR1-ACP1-M3-TE | Infusion | 152_V | P-MK478 | GATGGCGATCGGGTCATCGACGGGGGCCGTGG | pMK142 |
| | | | P-MK153 | AGCGGGACTCCCGCCC | |
| | | 152_I | P-MK398 | GACCCGATCGCCATCGTC | pRSG34 |
| | | | P-MK399 | GTCGAGCTGACTAGTGTGCTG | |
| pMK168 - SZ4-KR5-ACP5(2) | Infusion | 168_V | P-MK402 | GACGAGCCGCTCCAGGTA | pMK137 |
| | | | P-MK291 | TCGAGCTCCGTCGACAAGCT | |
| | | 168_I | P-MK457 | CTGGAGCGGCTCGTCGGCACCGAGGTCCGGG | pMK112 |
| | | | P-MK458 | GTCGACGGAGCTCGAATTCGGATCGCCGTCGAGC | |
| pMK169 - SZ4-RIFS(KR1-ACP1)(2) | Infusion | 169_V | P-MK459 | GAAGAGCTTGGCGCGCAG | pMK138 |
| | | | P-MK291 | TCGAGCTCCGTCGACAAGCT | |
| | | 169_I | P-MK460 | CGCGCCAAGCTCTTCGGCACCGAGGTCCGGG | pMK112 |
| | | | P-MK458 | GTCGACGGAGCTCGAATTCGGATCGCCGTCGAGC | |
| pMK170 - SZ4-M5-TE | Infusion | 170_V | digestion of pMK146 with HindIII | | |
| | | 170_I | P-MK390 | CGGATCCGGTAAGCTTGAGCCGATCGCGATCGTC | pBL17 |
| | | | P-MK389 | GTGCGGCCGCAAGCTTCGAATTCCCTCCGCCCAG | |

References for plasmids constructed elsewhere: pCK7[1]; pBP130[2]; pAYC59[3]; pAJ20 and pAJ21[4]; pBL13, pBL16, pBL17,and pBL18[5]; pRSG34[6]; pfRSZ was a gift from Mislav Oreb, Goethe University Frankfurt. In addition pBL12 was used to produce LDD(4)[5].

**Table S2: Amino acid sequences of proteins used in this study**. If no indication is given the PKS domain/modules are derived from DEBS. SZ domains are shown in red, docking domains in green, and linker regions in gray. $L_{12}$ is the ACP1-KS2 linker used to covalently fuse ACP1 to a heterologous acceptor module.

| Construct | Amino acid sequence |
|---|---|
| MK96<br>KR1-ACP1-(2) | MDEVSALRYRIEWRPTGAGEPARLDGTWLVAKYAGTADETSTAAREALESAGARVRELVVDARCGRDELAERLRSVGEV<br>AGVLSLLAVDEAEPEEAPLALASLADTLSLVQAMVSAELGCPLWTVTESAVATGPFERVRNAAHGALWGVGRVIALENPA<br>VWGGLVDVPAGSVAELARHLAAVVSGGAGEDQLALRADGVYGRRWVRAAAPATDDEWKPTGTVLVTGGTGGVGGQIA<br>RWLARRGAPHLLLVSRSGPDADGAGELVAELEALGARTTVAACDVTDRESVRELLGGIGDDVPLSAVFHAAATLDDGTVD<br>TLTGERIERASRAKVLGARNLHELTRELDLTAFVLFSSFASAFGAPGLGGYAPGNAYLDGLAQQRRSDGLPATAVAWGTW<br>AGSGMAEGPVADRFRRHGVIEMPPETACRALQNALDRAEVCPIVIDVRWDRFLLAYTAQRPTRLFDEIDDARRAAPQAAAE<br>PRVGALASLPAPEREKALFELVRSHAAAVLGHASAERVPADQAFAELGVDSLSALELRNRLGAATGVRLPTTTVFDHPDVR<br>TLAAHLAAELGTEVRGEAPSALAGLDALEAALPEVPATEREELVQRLERMLAALRPVAQAADASGTGANPSGDDLGEAGV<br>DELLEALGRELDGDPNSSSVDKLAAALEHHHHHH* |
| MK111<br>(5)KS1-AT1-L-SZ3 | MSGDNGMTEEKLRRYLKRTVTELDSVTARLREVEHRAGEPVAVVAMACRLPGGVSTPEEFWELLSEGRDAVAGLPTDRG<br>WDLDSLFHPDPTRSGTAHQRGGGFLTEATAFDPAFFGMSPREALAVDPQQRLMLELSWEVLERAGIPPTSLQASPTGVFVGL<br>IPQEYGPRLAEGGEGVEGYLMTGTTTSVASGRIAYTLGLEGPAISVDTACSSSLVAVHLACQSLRRGESSLAMAGGVTVMPT<br>PGMLVDFSRMNSLAPDGRCKAFSAGANGFGMAEGAGMLLLERLSDARRNGHPVLAVLRGTAVNSDGASNGLSAPNGRAQ<br>VRVIQQALAESGLGPADIDAVEAHGTGTRLGDPIEARALFEAYGRDREQPLHLGSVKSNLGHTQAAAGVAGVIKMVLAMR<br>AGTLPRTLHASERSKEIDWSSGAISLLDEPEPWPAGARPRRAGVSSFGISGTNAHAIIEEAPQVVEGERVEAGDVVAPWVLSA<br>SSAEGLRAQAARLAAHLREHPGQDPRDIAYSLATGRAALPHRAAFAPVDESAALRVLDGLATGNADGAAVGTSRAQQRAV<br>FVFPGQGWQWAGMAVDLLDTSPVFAAALRECADALEPHLDFEVIPFLRAEAARREQDAALSTERVDVVQPVMFAVMVSL<br>ASMWRAHGVEPAAVIGHSQGEIAAACVAGALSLDDAARVVALRSRVIATMPGNKGMASIAAPAGEVRARIGDRVEIAAVN<br>GPRSVVVAGDSDELDRLVASCTTECIRAKRLAVDYASHSSHVETIRDALHAELGEDFHPLPGFVPFFSTVTGRWTQPDELDA<br>GYWYRNLRRTVRFADAVRALAEQGYRTFLEVSAHPILTAAIEEIGDGSGADLSAIHSLRRGDGSLADFGEALSRAFAAGVA<br>VDWESVHLGTGARRVPLPTYPFQRERVWLGGSGGGSGNEVTTLENDAAFIENENAYLEKEIARLRKEKAALRNRLAHKKL<br>EHHHHHH* |
| MK112<br>SZ4-L-KR1-ACP1(2) | MQKVAELKNRVAVKLNRNEQLKNKVEELKNRNAYLKNELATLENEVARLENDVAEGGSGGGSGDEVSALRYRIEWRPTG<br>AGEPARLDGTWLVAKYAGTADETSTAAREALESAGARVRELVVDARCGRDELAERLRSVGEVAGVLSLLAVDEAEPEEAP<br>LALASLADTLSLVQAMVSAELGCPLWTVTESAVATGPFERVRNAAHGALWGVGRVIALENPAVWGGLVDVPAGSVAELA<br>RHLAAVVSGGAGEDQLALRADGVYGRRWVRAAAPATDDEWKPTGTVLVTGGTGGVGGQIARWLARRGAPHLLLVSRSG<br>PDADGAGELVAELEALGARTTVAACDVTDRESVRELLGGIGDDVPLSAVFHAAATLDDGTVDTLTGERIERASRAKVLGA<br>RNLHELTRELDLTAFVLFSSFASAFGAPGLGGYAPGNAYLDGLAQQRRSDGLPATAVAWGTWAGSGMAEGPVADRFRRH<br>GVIEMPPETACRALQNALDRAEVCPIVIDVRWDRFLLAYTAQRPTRLFDEIDDARRAAPQAAAEPRVGALASLPAPEREKAL<br>FELVRSHAAAVLGHASAERVPADQAFAELGVDSLSALELRNRLGAATGVRLPTTTVFDHPDVRTLAAHLAAELGTEVRGE<br>APSALAGLDALEAALPEVPATEREELVQRLERMLAALRPVAQAADASGTGANPSGDDLGEAGVDELLEALGRELDGDPNS<br>SSVDKLAAALEHHHHHH* |
| MK113<br>(5)KS1-AT1 | MSGDNGMTEEKLRRYLKRTVTELDSVTARLREVEHRAGEPVAVVAMACRLPGGVSTPEEFWELLSEGRDAVAGLPTDRG<br>WDLDSLFHPDPTRSGTAHQRGGGFLTEATAFDPAFFGMSPREALAVDPQQRLMLELSWEVLERAGIPPTSLQASPTGVFVGL<br>IPQEYGPRLAEGGEGVEGYLMTGTTTSVASGRIAYTLGLEGPAISVDTACSSSLVAVHLACQSLRRGESSLAMAGGVTVMPT<br>PGMLVDFSRMNSLAPDGRCKAFSAGANGFGMAEGAGMLLLERLSDARRNGHPVLAVLRGTAVNSDGASNGLSAPNGRAQ |

| | |
|---|---|
| | VRVIQQALAESGLGPADIDAVEAHGTGTRLGDPIEARALFEAYGRDREQPLHLGSVKSNLGHTQAAAGVAGVIKMVLAMR<br>AGTLPRTLHASERSKEIDWSSGAISLLDEPEPWPAGARPRRAGVSSFGISGTNAHAIIEEAPQVVEGERVEAGDVVAPWVLSA<br>SSAEGLRAQAARLAAHLREHPGQDPRDIAYSLATGRAALPHRAAFAPVDESAALRVLDGLATGNADGAAVGTSRAQQRAV<br>FVFPGQGWQWAGMAVDLLDTSPVFAAALRECADALEPHLDFEVIPFLRAEAARREQDAALSTERVDVVQPVMFAVMVSL<br>ASMWRAHGVEPAAVIGHSQGEIAAACVAGALSLDDAARVVALRSRVIATMPGNKGMASIAAPAGEVRARIGDRVEIAAVN<br>GPRSVVVAGDSDELDRLVASCTTECIRAKRLAVDYASHSSHVETIRDALHAELGEDFHPLPGFVPFFSTVTGRWTQPDELDA<br>GYWYRNLRRTVRFADAVRALAEQGYRTFLEVSAHPILTAAIEEIGDGSGADLSAIHSLRRGDGSLADFGEALSRAFAAGVA<br>VDWESVHLGTGARRVPLPTYPFQRERVWLLEHHHHHH* |
| MK136<br>SZ4-L-KR1-ACP1-<br>$L_{12}$-M2-TE | MQKVAELKNRVAVKLNRNEQLKNKVEELKNRNAYLKNELATLENEVARLENDVAEGGSGGGSGDEVSALRYRIEWRPTG<br>AGEPARLDGTWLVAKYAGTADETSTAAREALESAGARVRELVVDARCGRDELAERLRSVGEVAGVLSLLAVDEAEPEEAP<br>LALASLADTLSLVQAMVSAELGCPLWTVTESAVATGPFERVRNAAHGALWGVGRVIALENPAVWGGLVDVPAGSVAELA<br>RHLAAVVSGGAGEDQLALRADGVYGRRWVRAAAPATDDEWKPTGTVLVTGGTGGVGGQIARWLARRGAPHLLLVSRSG<br>PDADGAGELVAELEALGARTTVAACDVTDRESVRELLGGIGDDVPLSAVFHAAATLDDGTVDTLTGERIERASRAKVLGA<br>RNLHELTRELDLTAFVLFSSFASAFGAPGLGGYAPGNAYLDGLAQQRRSDGLPATAVAWGTWAGSGMAEGPVADRFRRH<br>GVIEMPPETACRALQNALDRAEVCPIVIDVRWDRFLLAYTAQRPTRLFDEIDDARRAAPQAAAEPRVGAHMLASLPAPERE<br>KALFELVRSHAAAVLGHASAERVPADQAFAELGVDSLSALELRNRLGAATGVRLPTTTVFDHPDVRTLAAHLAAELGGAT<br>GAEQAAPATTAPVDEPIAIVGMACRLPGEVDSPERLWELITSGRDSAAEVPDDRGWVPDELMASDAAGTRRAHGNFMAGA<br>GDFDAAFFGISPREALAMDPQQRQALETTWEALESAGIPPETLRGSDTGVFVGMSHQGYATGRPRPEDGVDGYLLTGNTAS<br>VASGRIAYVLGLEGPALTVDTACSSSLVALHTACGSLRDGDCGLAVAGGVSVMAGPEVFTEFSRQGALSPDGRCKPFSDEA<br>DGFGLGEGSAFVVLQRLSDARREGRRVLGVVAGSAVNQDGASNGLSAPSGVAQQRVIRRAWARAGITGADVAVVEAHGT<br>GTRLGDPVEASALLATYGKSRGSSGPVLLGSVKSNIGHAQAAAGVAGVIKVLLGLERGVVPPMLCRGERSGLIDWSSGEIEL<br>ADGVREWSPAADGVRRAGVSAFGVSGTNAHVIIAEPPEPEPVPQPRRMLPATGVVPVVLSARTGAALRAQAGRLADHLAA<br>HPGIAPADVSWTMARARQHFEERAAVLAADTAEAVHRLRAVADGAVVPGVVTGSASDGGSVFVFPGQGAQWEGMAREL<br>LPVPVFAESIAECDAVLSEVAGFSVSEVLEPRPDAPSLERVDVVQPVLFAVMVSLARLWRACGAVPSAVIGHSQGEIAAAVV<br>AGALSLEDGMRVVARRSRAVRAVAGRGSMLSVRGGRSDVEKLLADDSWTGRLEVAAVNGPDAVVVAGDAQAAREFLEY<br>CEGVGIRARAIPVDYASHTAHVEPVRDELVQALAGITPRRAEVPFFSTLTGDFLDGTELDAGYWYRNLRHPVEFHSAVQAL<br>TDQGYATFIEVSPHPVLASSVQETLDDAESDAAVLGTLERDAGDADRFLTALADAHTRGVAVDWEAVLGRAGLVDLPGYP<br>FQGKRFWLLPDRTTPRDELDGWFYRVDWTEVPRSEPAALRGRWLVVVPEGHEEDGWTVEVRSALAEAGAEPEVTRGVGG<br>LVGDCAGVVSLLALEGDGAVQTLVLVRELDAEGIDAPLWTVTFGAVDAGSPVARPDQAKLWGLGQVASLERGPRWTGLV<br>DLPHMPDPELRGRLTAVLAGSEDQVAVRADAVRARRLSPAHVTATSEYAVPGGTILVTGGTAGLGAEVARWLAGRGAEH<br>LALVSRRGPDTEGVGDLTAELTRLGARVSVHACDVSSREPVRELVHGLIEQGDVVRGVVHAAGLPQQVAINDMDEAAFDE<br>VVAAKAGGAVHLDELCSDAELFLLFSSGAGVWGSARQGAYAAGNAFLDAFARHRRGRGLPATSVAWGLWAAGGMTGD<br>EEAVSFLRERGVRAMPVPRALAALDRVLASGETAVVVTDVDWPAFAESYTAARPRPLLDRIVTTAPSERAGEPETESLRDR<br>LAGLPRAERTAELVRLVRTSTATVLGHDDPKAVRATTPFKELGFDSLAAVRLRNLLNAATGLRLPSTLVFDHPNASAVAGF<br>LTSELGSGTPAREASSALRDGYRQAGVSGRVRSYLDLLAGLSDFREHFDGSDGFSLDLVDMADGPGEVTVICCAGTAAISGP<br>HEFTRLAGALRGIAPVRAVPQPGYEEGEPLPSSMAAVAAVQADAVIRTQGDKPFVVAGHSAGALMAYALATELLDRGHPP<br>RGVVLIDVYPPGHQDAMNAWLEELTATLFDRETVRMDDTRLTALGAYDRLTGQWRPRETGLPTLLVSAGEPMGPWPDDS<br>WKPTWPFEHDTVAVPGDHFTMVQEHADAIARHIDAWLGGGNSSSVDKLAAALEHHHHHH* |

| | |
|---|---|
| MK137<br>SZ4-L-KR5-ACP5-<br>M6-TE | MQKVAELKNRVAVKLNRNEQLKNKVEELKNRNAYLKNELATLENEVARLENDVAEGGSGGGSGPIPTGGRARDEDDDW<br>RYQVVVWREAEWESASLAGRVLLVTGPGVPSELSDAIRSGLEQSGATVLTCDVESRSTIGTALEAADTDALSTVVSLLSRDGE<br>AVDPSLDALALVQALGAAGVEAPLWVLTRNAVQVADGELVDPAQAMVGGLGRVVGIEQPGRWGGLVDLVDADAASIRS<br>LAAVLADPRGEEQVAIRADGIKVARLVPAPARAARTRWSPRGTVLVTGGTGGIGAHVARWLARSGAEHLVLLGRRGADAP<br>GASELREELTALGTGVTIAACDVADRARLEAVLAAERAEGRTVSAVMHAAGVSTSTPLDDLTEAEFTEIADVKVRGTVNLD<br>ELCPDLDAFVLFSSNAGVWGSPGLASYAAANAFLDGFARRRRSEGAPVTSIAWGLWAGQNMAGDEGGEYLRSQGLRAMD<br>PDRAVEELHITLDHGQTSVSVVDMDRRRFVELFTAARHRPLFDEIAGARAEARQSEEGPALAQRLAALSTAERREHLAHLIR<br>AEVAAVLGHGDDAAIDRDRAFRDLGFDSMTAVDLRNRLAAVTGVREAATVVFDHPTITRLADHYLERLVGAAEAEQAPA<br>LVREVPKDADDPIAIVGMACRFPGGVHNPGELWEFIVGGGDAVTEMPTDRGWDLDALFDPDPQRHGTSYSRHGAFLDGAA<br>DFDAAFFGISPREALAMDPQQRQVLETTWELFENAGIDPHSLRGSDTGVFLGAAYQGYGQDAVVPEDSEGYLLTGNSSAVV<br>SGRVAYVLGLEGPAVTVDTACSSSLVALHSACGSLRDGDCGLAVAGGVSVMAGPEVFTEFSRQGGLAVDGRCKAFSAEAD<br>GFGFAEGVAVVLLQRLSDARRAGRQVLGVVAGSAINQDGASNGLAAPSGVAQQRVIRKAWARAGITGADVAVVEAHGTG<br>TRLGDPVEASALLATYGKSRGSSGPVLLGSVKSNIGHAQAAAGVAGVIKVVLGLNRGLVPPMLCRGERSPLIEWSSGGVEL<br>AEAVSPWPPAADGVRRAGVSAFGVSGTNAHVIIAEPPEPEPLPEPGPVGVLAAANSVPVLLSARTETALAAQARLLESAVDD<br>SVPLTALASALATGRAHLPRRAALLAGDHEQLRGQLRAVAEGVAAPGATTGTASAGGVVFVFPGQGAQWEGMARGLLSV<br>PVFAESIAECDAVLSEVAGFSASEVLEQRPDAPSLERVDVVQPVLFSVMVSLARLWGACGVSPSAVIGHSQGEIAAAVVAG<br>VLSLEDGVRVVALRAKALRALAGKGGMVSLAAPGERARALIAPWEDRISVAAVNSPSSVVVSGDPEALAELVARCEDEGV<br>RAKTLPVDYASHSRHVEEIRETILADLDGISARRAAIPLYSTLHGERRDGADMGPRYWYDNLRSQVRFDEAVSAAVADGHA<br>TFVEMSPHPVLTAAVQEIAADAVAIGSLHRDTAEEHLIAELARAHVHGVAVDWRNVFPAAPPVALPNYPFEPQRYWLAPEV<br>SDQLADSRYRVDWRPLATTPVDLEGGFLVHGSAPESLTSAVEKAGGRVVPVASADREALAAALREVPGEVAGVLSVHTGA<br>ATHLALHQSLGEAGVRAPLWLVTSRAVALGESEPVDPEQAMVWGLGRVMGLETPERWGGLVDLPAEPAPGDGEAFVACL<br>GADGHEDQVAIRDHARYGRRLVRAPLGTRESSWEPAGTALVTGGTGALGGHVARHLARCGVEDLVLVSRRGVDAPGAAE<br>LEAELVALGAKTTITACDVADREQLSKLLEELRGQGRPVRTVVHTAGVPESRPLHEIGELESVCAAKVTGARLLDELCPDAE<br>TFVLFSSGAGVWGSANLGAYSAANAYLDALAHRRRAEGRAATSVAWGAWAGEGMATGDLEGLTRRGLRPMAPERAIRA<br>LHQALDNGDTCVSIADVDWERFAVGFTAARPRPLLDELVTPAVGAVPAVQAAPAREMTSQELLEFTHSHVAAILGHSSPDA<br>VGQDQPFTELGFDSLTAVGLRNQLQQATGLALPATLVFEHPTVRRLADHIGQQLDSGTPAREASSALRDGYRQAGVSGRVR<br>SYLDLLAGLSDFREHFDGSDGFSLDLVDMADGPGEVTVICCAGTAAISGPHEFTRLAGALRGIAPVRAVPQPGYEEGEPLPSS<br>MAAVAAVQADAVIRTQGDKPFVVAGHSAGALMAYALATELLDRGHPPRGVVLIDVYPPGHQDAMNAWLEELTATLFDRE<br>TVRMDDTRLTALGAYDRLTGQWRPRETGLPTLLVSAGEPMGPWPDDSWKPTWPFEHDTVAVPGDHFTMVQEHADAIARH<br>IDAWLGGGNSSSVDKLAAALEHHHHHH* |
| MK138<br>SZ4-L-RIFS (KR1-<br>ACP1-M2)-TE | MQKVAELKNRVAVKLNRNEQLKNKVEELKNRNAYLKNELATLENEVARLENDVAEGGSGGGSGEPAEPASAGDPLLGTV<br>VSTPGSDRLTAVAQWSRRAQPWAVDGLVPNAALVEAAIRLGDLAGTPVVGELVVDAPVVLPRRGSREVQLIVGEPGEQRR<br>RPIEVFSREADEPWTRHAHGTLAPAAAAVPEPAAAGDATDVTVAGLRDADRYGIHPALLDAAVRTVVGDDLLPSVWTGVS<br>LLASGATAVTVTPTATGLRLTDPAGQPVLTVESVRGTPFVAEQGTTDALFRVDWPEIPLPTAETADFLPYEATSAEATLSAL<br>QAWLADPAETRLAVVTGDCTEPGAAAIWGLVRSAQSEHPGRIVLADLDDPAVLPAVVASGEPQVRVRNGVASVPRLTRVT<br>PRQDARPLDPEGTVLITGGTGTLGALTARHLVTAHGVRHLVLVSRRGEAPELQEELTALGASVAIAACDVADRAQLEAVLR<br>AIPAEHPLTAVIHTAGVLDDGVVTELTPDRLATVRRPKVDAARLLDELTREADLAAFVLFSSAAGVLGNPGQAGYAAANAE<br>LDALARQRNSLDLPAVSIAWGYWATVSGMTEHLGDADLRRNQRIGMSGLPADEGMALLDAAIATGGTLVAAKFDVAALR<br>ATAKAGGPVPPLLRGLAPLPRRAAAKTASLTERLAGLAETEQAAALLDLVRRHAAEVLGHSGAESVHSGRTFKDAGFDSLT<br>AVELRNRLAAATGLTLSPAMIFDYPKPPALADHLRAKLFGSAANRPAEIGTAAAEEPIAIVAMACRFPGGVHSPEDLWRLVA |

| | |
|---|---|
| | DGADAVTEFPADRGWDTDRLYHEDPDHEGTTYVRHGAFLDDAAGFDAAFFGISPNEALAMDPQQRLLLETSWELFERAAI<br>DPTTLAGQDIGVFAGVNSHDYSMRMHRAAGVEGFRLTGGSASVLSGRVAYHFGVEGPAVTVDTACSSSLVALHMAVQAL<br>QRGECSMALAGGVMVMGTVETFVEFSRQRGLAPDGRCKAFADGADGTGWSEGVGLLLVERLSEAQRRGHQVLAVVRGS<br>AVNSDGASNGLTAPNGPSQQRVIRKALAAAGLSTSDVDAVEAHGTGTTLGDPIEAEALLATYGQNRETPLWLGSVKSNLG<br>HTQAAAGVAGVIKMVMAMRHGVLPRTLHVDRPSSYVDWSAGAVELLTEARDWVSNGHPRRAGVSSFGIGGTNAHVVLE<br>EVAAPITTPQPEPAEFLVPVLVSARTAAGLRGQAGRLAAFLGDRTDVRVPDAAYALATTRAQLDHRAVVLASDRAQLCAD<br>LAAFGSGVVTGTPVDGKLAVLFTGQGSQWAGMGRELAETFPVFRDAFEAACEAVDTHLRERPLREVVFDDSALLDQTMYT<br>QGALFAVETALFRLFESWGVRPGLLAGHSIGELAAAHVSGVLDLADAGELVAARGRLMQALPAGGAMVAVQATEDEVAP<br>LLDGTVCVAAVNGPDSVVLSGTEAAVLAVADELAGRGRKTRRLAVSHAFHSPLMEPMLDDFRAVAERLTYRAGSLPVVST<br>LTGELAALDSPDYWVGQVRNAVRFSDAVTALGAQGASTFLELGPGGALAAMALGTLGGPEQSCVATLRKNGAEVPDVLT<br>ALAELHVRGVGVDWTTVLDEPATAVGTVLPTYAFQHQRFWVDVDETAAVSVTPPPAEPIVDRPVQDVLELVRESAAVVLG<br>HRDAGSFDLDRSFKDHGFDSLSAVKLRNRLRDFTGVELPSTLIFDYPNPAVLADHLRAELLSGTPAREASSALRDGYRQAGV<br>SGRVRSYLDLLAGLSDFREHFDGSDGFSLDLVDMADGPGEVTVICCAGTAAISGPHEFTRLAGALRGIAPVRAVPQPGYEEG<br>EPLPSSMAAVAAVQADAVIRTQGDKPFVVAGHSAGALMAYALATELLDRGHPPRGVVLIDVYPPGHQDAMNAWLEELTA<br>TLFDRETVRMDDTRLTALGAYDRLTGQWRPRETGLPTLLVSAGEPMGPWPDDSWKPTWPFEHDTVAVPGDHFTMVQEHA<br>DAIARHIDAWLGGGNSSSVDKLAAALEHHHHHH* |
| MK141<br>SZ4-L-KR1-ACP1-<br>$L_{12}$-M5-TE | MQKVAELKNRVAVKLNRNEQLKNKVEELKNRNAYLKNELATLENEVARLENDVAEGGSGGGSGDEVSALRYRIEWRPTG<br>AGEPARLDGTWLVAKYAGTADETSTAAREALESAGARVRELVVDARCGRDELAERLRSVGEVAGVLSLLAVDEAEPEEAP<br>LALASLADTLSLVQAMVSAELGCPLWTVTESAVATGPFERVRNAAHGALWGVGRVIALENPAVWGGLVDVPAGSVAELA<br>RHLAAVVSGGAGEDQLALRADGVYGRRWVRAAAPATDDEWKPTGTVLVTGGTGGVGGQIARWLARRGAPHLLLVSRSG<br>PDADGAGELVAELEALGARTTVAACDVTDRESVRELLGGIGDDVPLSAVFHAAATLDDGTVDTLTGERIERASRAKVLGA<br>RNLHELTRELDLTAFVLFSSFASAFGAPGLGGYAPGNAYLDGLAQQRRSDGLPATAVAWGTWAGSGMAEGPVADRFRRH<br>GVIEMPPETACRALQNALDRAEVCPIVIDVRWDRFLLAYTAQRPTRLFDEIDDARRAAPQAAAEPRVGAHMLASLPAPERE<br>KALFELVRSHAAAVLGHASAERVPADQAFAELGVDSLSALELRNRLGAATGVRLPTTTVFDHPDVRTLAAHLAAELGGAT<br>GAEQAAPATTAPVDEPIAIVGMACRFPGDVDSPESFWEFVSGGGDAIAEAPADRGWEPDPDARLGGMLAAAGDFDAGFFGI<br>SPREALAMDPQQRIMLEISWEALERAGHDPVSLRGSATGVFTGVGTVDYGPRPDEAPDEVLGYVGTGTASSVASGRVAYCL<br>GLEGPAMTVDTACSSGLTALHLAMESLRRDECGLALAGGVTVMSSPGAFTEFRSQGGLAADGRCKPFSKAADGFGLAEGA<br>GVLVLQRLSAARREGRPVLAVLRGSAVNQDGASNGLTAPSGPAQQRVIRRALENAGVRAGDVDYVEAHGTGTRLGDPIEV<br>HALLSTYGAERDPDDPLWIGSVKSNIGHTQAAAGVAGVMKAVLALRHGEMPRTLHFDEPSPQIEWDLGAVSVVSQARSWP<br>AGERPRRAGVSSFGISGTNAHVIVEEAPEADEPEPAPDSGPVPLVLSGRDEQAMRAQAGRLADHLAREPRNSLRDTGFTLAT<br>RRSAWEHRAVVVGDRDDALAGLRAVADGRIADRTATGQARTRRGVAMVFPGQGAQWQGMARDLLRESQVFADSIRDCE<br>RALAPHVDWSLTDLLSGARPLDRVDVVQPALFAVMVSLAALWRSHGVEPAAVVGHSQGEIAAAHVAGALTLEDAAKLVA<br>VRSRVLRRLGGQGGMASFGLGTEQAAERIGRFAGALSIASVNGPRSVVVAGESGPLDELIAECEAEGITARRIPVDYASHSPQ<br>VESLREELLTELAGISPVSADVALYSTTTGQPIDTATMDTAYWYANLREQVRFQDATRQLAEAGFDAFVEVSPHPVLTVGIE<br>ATLDSALPADAGACVVGTLRRDRGGLADFHTALGEAYAQGVEVDWSPAFADARPVELPVYPFQRQRYWLPIPTGGRARDE<br>DDDWRYQVVWREAEWESASLAGRVLLVTGPGVPSELSDAIRSGLEQSGATVLTCDVESRSTIGTALEAADTDALSTVVSLL<br>SRDGEAVDPSLDALALVQALGAAGVEAPLWVLTRNAVQVADGELVDPAQAMVGGLGRVVGIEQPGRWGGLVDLVDADA<br>ASIRSLAAVLADPRGEEQVAIRADGIKVARLVPAPARAARTRWSPRGTVLVTGGTGGIGAHVARWLARSGAEHLVLLGRR<br>GADAPGASELREELTALGTGVTIAACDVADRARLEAVLAAERAEGRTVSAVMHAAGVSTSTPLDDLTEAEFTEIADVKVRG<br>TVNLDELCPDLDAFVLFSSNAGVWGSPGLASYAAANAFLDGFARRRRSEGAPVTSIAWGLWAGQNMAGDEGGEYLRSQG |

| | |
|---|---|
| | LRAMDPDRAVEELHITLDHGQTSVSVVDMDRRRFVELFTAARHRPLFDEIAGARAEARQSEEGPALAQRLAALSTAERREH<br>LAHLIRAEVAAVLGHGDDAAIDRDRAFRDLGFDSMTAVDLRNRLAAVTGVREAATVVFDHPTITRLADHYLERLVSGTPA<br>REASSALRDGYRQAGVSGRVRSYLDLLAGLSDFREHFDGSDGFSLDLVDMADGPGEVTVICCAGTAAISGPHEFTRLAGAL<br>RGIAPVRAVPQPGYEEGEPLPSSMAAVAAVQADAVIRTQGDKPFVVAGHSAGALMAYALATELLDRGHPPRGVVLIDVYP<br>PGHQDAMNAWLEELTATLFDRETVRMDDTRLTALGAYDRLTGQWRPRETGLPTLLVSAGEPMGPWPDDSWKPTWPFEHD<br>TVAVPGDHFTMVQEHADAIARHIDAWLGGGNSSSVDKLAAALEHHHHHH* |
| MK142<br>SZ4-L-KR1-ACP1-<br>$L_{12}$-M6-TE | MQKVAELKNRVAVKLNRNEQLKNKVEELKNRNAYLKNELATLENEVARLENDVAEGGSGGGSGDEVSALRYRIEWRPTG<br>AGEPARLDGTWLVAKYAGTADETSTAAREALESAGARVRELVVDARCGRDELAERLRSVGEVAGVLSLLAVDEAEPEEAP<br>LALASLADTLSLVQAMVSAELGCPLWTVTESAVATGPFERVRNAAHGALWGVGRVIALENPAVWGGLVDVPAGSVAELA<br>RHLAAVVSGGAGEDQLALRADGVYGRRWVRAAAPATDDEWKPTGTVLVTGGTGGVGGQIARWLARRGAPHLLLVSRSG<br>PDADGAGELVAELEALGARTTVAACDVTDRESVRELLGGIGDDVPLSAVFHAAATLDDGTVDTLTGERIERASRAKVLGA<br>RNLHELTRELDLTAFVLFSSFASAFGAPGLGGYAPGNAYLDGLAQQRRSDGLPATAVAWGTWAGSGMAEGPVADRFRRH<br>GVIEMPPETACRALQNALDRAEVCPIVIDVRWDRFLLAYTAQRPTRLFDEIDDARRAAPQAAAEPRVGAHMLASLPAPERE<br>KALFELVRSHAAAVLGHASAERVPADQAFAELGVDSLSALELRNRLGAATGVRLPTTTVFDHPDVRTLAAHLAAELGGAT<br>GAEQAAPATTAPVDDPIAIVGMACRFPGGVHNPGELWEFIVGGGDAVTEMPTDRGWDLDALFDPDPQRHGTSYSRHGAFL<br>DGAADFDAAFFGISPREALAMDPQQRQVLETTWELFENAGIDPHSLRGSDTGVFLGAAYQGYGQDAVVPEDSEGYLLTGN<br>SSAVVSGRVAYVLGLEGPAVTVDTACSSSLVALHSACGSLRDGDCGLAVAGGVSVMAGPEVFTEFSRQGGLAVDGRCKAF<br>SAEADGFGFAEGVAVVLLQRLSDARRAGRQVLGVVAGSAINQDGASNGLAAPSGVAQQRVIRKAWARAGITGADVAVVE<br>AHGTGTRLGDPVEASALLATYGKSRGSSGPVLLGSVKSNIGHAQAAAGVAGVIKVVLGLNRGLVPPMLCRGERSPLIEWSS<br>GGVELAEAVSPWPPAADGVRRAGVSAFGVSGTNAHVIIAEPPEPEPLPEPGPVGVLAAANSVPVLLSARTETALAAQARLLE<br>SAVDDSVPLTALASALATGRAHLPRRAALLAGDHEQLRGQLRAVAEGVAAPGATTGTASAGGVVFVFPGQGAQWEGMAR<br>GLLSVPVFAESIAECDAVLSEVAGFSASEVLEQRPDAPSLERVDVVQPVLFSVMVSLARLWGACGVSPSAVIGHSQGEIAAA<br>VVAGVLSLEDGVRVVALRAKALRALAGKGGMVSLAAPGERARALIAPWEDRISVAAVNSPSSVVVSGDPEALAELVARCE<br>DEGVRAKTLPVDYASHSRHVEEIRETILADLDGISARRAAIPLYSTLHGERRDGADMGPRYWYDNLRSQVRFDEAVSAAVA<br>DGHATFVEMSPHPVLTAAVQEIAADAVAIGSLHRDTAEEHLIAELARAHVHGVAVDWRNVFPAAPPVALPNYPFEPQRYW<br>LAPEVSDQLADSRYRVDWRPLATTPVDLEGGFLVHGSAPESLTSAVEKAGGRVVPVASADREALAAALREVPGEVAGVLS<br>VHTGAATHLALHQSLGEAGVRAPLWLVTSRAVALGESEPVDPEQAMVWGLGRVMGLETPERWGGLVDLPAEPAPGDGE<br>AFVACLGADGHEDQVAIRDHARYGRRLVRAPLGTRESSWEPAGTALVTGGTGALGGHVARHLARCGVEDLVLVSRRGVD<br>APGAAELEAELVALGAKTTITACDVADREQLSKLLEELRGQGRPVRTVVHTAGVPESRPLHEIGELESVCAAKVTGARLLD<br>ELCPDAETFVLFSSGAGVWGSANLGAYSAANAYLDALAHRRRAEGRAATSVAWGAWAGEGMATGDLEGLTRRGLRPMA<br>PERAIRALHQALDNGDTCVSIADVDWERFAVGFTAARPRPLLDELVTPAVGAVPAVQAAPAREMTSQELLEFTHSHVAAIL<br>GHSSPDAVGQDQPFTELGFDSLTAVGLRNQLQQATGLALPATLVFEHPTVRRLADHIGQQLDSGTPAREASSALRDGYRQA<br>GVSGRVRSYLDLLAGLSDFREHFDGSDGFSLDLVDMADGPGEVTVICCAGTAAISGPHEFTRLAGALRGIAPVRAVPQPGYE<br>EGEPLPSSMAAVAAVQADAVIRTQGDKPFVVAGHSAGALMAYALATELLDRGHPPRGVVLIDVYPPGHQDAMNAWLEEL<br>TATLFDRETVRMDDTRLTALGAYDRLTGQWRPRETGLPTLLVSAGEPMGPWPDDSWKPTWPFEHDTVAVPGDHFTMVQE<br>HADAIARHIDAWLGGGNSSSVDKLAAALEHHHHHH* |
| MK147<br>SZ4-L-M6-TE | MQKVAELKNRVAVKLNRNEQLKNKVEELKNRNAYLKNELATLENEVARLENDVAEGGSGGGSGKLDPIAIVGMACRFPG<br>GVHNPGELWEFIVGGGDAVTEMPTDRGWDLDALFDPDPQRHGTSYSRHGAFLDGAADFDAAFFGISPREALAMDPQQRQV<br>LETTWELFENAGIDPHSLRGSDTGVFLGAAYQGYGQDAVVPEDSEGYLLTGNSSAVVSGRVAYVLGLEGPAVTVDTACSSS<br>LVALHSACGSLRDGDCGLAVAGGVSVMAGPEVFTEFSRQGGLAVDGRCKAFSAEADGFGFAEGVAVVLLQRLSDARRAG |

| | |
|---|---|
| | RQVLGVVAGSAINQDGASNGLAAPSGVAQQRVIRKAWARAGITGADVAVVEAHGTGTRLGDPVEASALLATYGKSRGSS<br>GPVLLGSVKSNIGHAQAAAGVAGVIKVVLGLNRGLVPPMLCRGERSPLIEWSSGGVELAEAVSPWPPAADGVRRAGVSAF<br>GVSGTNAHVIIAEPPEPEPLPEPGPVGVLAAANSVPVLLSARTETALAAQARLLESAVDDSVPLTALASALATGRAHLPRRA<br>ALLAGDHEQLRGQLRAVAEGVAAPGATTGTASAGGVVFVFPGQGAQWEGMARGLLSVPVFAESIAECDAVLSEVAGFSAS<br>EVLEQRPDAPSLERVDVVQPVLFSVMVSLARLWGACGVSPSAVIGHSQGEIAAAVVAGVLSLEDGVRVVALRAKALRALA<br>GKGGMVSLAAPGERARALIAPWEDRISVAAVNSPSSVVVSGDPEALAELVARCEDEGVRAKTLPVDYASHSRHVEEIRETIL<br>ADLDGISARRAAIPLYSTLHGERRDGADMGPRYWYDNLRSQVRFDEAVSAAVADGHATFVEMSPHPVLTAAVQEIAADAV<br>AIGSLHRDTAEEHLIAELARAHVHGVAVDWRNVFPAAPPVALPNYPFEPQRYWLAPEVSDQLADSRYRVDWRPLATTPVD<br>LEGGFLVHGSAPESLTSAVEKAGGRVVPVASADREALAAALREVPGEVAGVLSVHTGAATHLALHQSLGEAGVRAPLWLV<br>TSRAVALGESEPVDPEQAMVWGLGRVMGLETPERWGGLVDLPAEPAPGDGEAFVACLGADGHEDQVAIRDHARYGRRLV<br>RAPLGTRESSWEPAGTALVTGGTGALGGHVARHLARCGVEDLVLVSRRGVDAPGAAELEAELVALGAKTTITACDVADRE<br>QLSKLLEELRGQGRPVRTVVHTAGVPESRPLHEIGELESVCAAKVTGARLLDELCPDAETFVLFSSGAGVWGSANLGAYSA<br>ANAYLDALAHRRRAEGRAATSVAWGAWAGEGMATGDLEGLTRRGLRPMAPERAIRALHQALDNGDTCVSIADVDWERF<br>AVGFTAARPRPLLDELVTPAVGAVPAVQAAPAREMTSQELLEFTHSHVAAILGHSSPDAVGQDQPFTELGFDSLTAVGLRN<br>QLQQATGLALPATLVFEHPTVRRLADHIGQQLDSGTPAREASSALRDGYRQAGVSGRVRSYLDLLAGLSDFREHFDGSDGF<br>SLDLVDMADGPGEVTVICCAGTAAISGPHEFTRLAGALRGIAPVRAVPQPGYEEGEPLPSSMAAVAAVQADAVIRTQGDKP<br>FVVAGHSAGALMAYALATELLDRGHPPRGVVLIDVYPPGHQDAMNAWLEELTATLFDRETVRMDDTRLTALGAYDRLTG<br>QWRPRETGLPTLLVSAGEPMGPWPDDSWKPTWPFEHDTVAVPGDHFTMVQEHADAIARHIDAWLGGGNSKLAAALEHHH<br>HHH* |
| MK148<br>SZ4-L-M2-TE | MQKVAELKNRVAVKLNRNEQLKNKVEELKNRNAYLKNELATLENEVARLENDVAEGGSGGGSGKLEPIAIVGMACRLPG<br>EVDSPERLWELITSGRDSAAEVPDDRGWVPDELMASDAAGTRRAHGNFMAGAGDFDAAFFGISPREALAMDPQQRQALET<br>TWEALESAGIPPETLRGSDTGVFVGMSHQGYATGRPRPEDGVDGYLLTGNTASVASGRIAYVLGLEGPALTVDTACSSSLV<br>ALHTACGSLRDGDCGLAVAGGVSVMAGPEVFTEFSRQGALSPDGRCKPFSDEADGFGLGEGSAFVVLQRLSDARREGRRV<br>LGVVAGSAVNQDGASNGLSAPSGVAQQRVIRRAWARAGITGADVAVVEAHGTGTRLGDPVEASALLATYGKSRGSSGPV<br>LLGSVKSNIGHAQAAAGVAGVIKVLLGLERGVVPPMLCRGERSGLIDWSSGEIELADGVREWSPAADGVRRAGVSAFGVS<br>GTNAHVIIAEPPEPEPVPQPRRMLPATGVVPVVLSARTGAALRAQAGRLADHLAAHPGIAPADVSWTMARARQHFEERAA<br>VLAADTAEAVHRLRAVADGAVVPGVVTGSASDGGSVFVFPGQGAQWEGMARELLPVPVFAESIAECDAVLSEVAGFSVSE<br>VLEPRPDAPSLERVDVVQPVLFAVMVSLARLWRACGAVPSAVIGHSQGEIAAAVVAGALSLEDGMRVVARRSRAVRAVA<br>GRGSMLSVRGGRSDVEKLLADDSWTGRLEVAAVNGPDAVVVAGDAQAAREFLEYCEGVGIRARAIPVDYASHTAHVEPV<br>RDELVQALAGITPRRAEVPFFSTLTGDFLDGTELDAGYWYRNLRHPVEFHSAVQALTDQGYATFIEVSPHPVLASSVQETLD<br>DAESDAAVLGTLERDAGDADRFLTALADAHTRGVAVDWEAVLGRAGLVDLPGYPFQGKRFWLLPDRTTPRDELDGWFYR<br>VDWTEVPRSEPAALRGRWLVVVPEGHEEDGWTVEVRSALAEAGAEPEVTRGVGGLVGDCAGVVSLLALEGDGAVQTLVL<br>VRELDAEGIDAPLWTVTFGAVDAGSPVARPDQAKLWGLGQVASLERGPRWTGLVDLPHMPDPELRGRLTAVLAGSEDQV<br>AVRADAVRARRLSPAHVTATSEYAVPGGTILVTGGTAGLGAEVARWLAGRGAEHLALVSRRGPDTEGVGDLTAELTRLGA<br>RVSVHACDVSSREPVRELVHGLIEQGDVVRGVVHAAGLPQQVAINDMDEAAFDEVVAAKAGGAVHLDELCSDAELFLLFS<br>SGAGVWGSARQGAYAAGNAFLDAFARHRRGRGLPATSVAWGLWAAGGMTGDEEAVSFLRERGVRAMPVPRALAALDR<br>VLASGETAVVVTDVDWPAFAESYTAARPRPLLDRIVTTAPSERAGEPETESLRDRLAGLPRAERTAELVRLVRTSTATVLGH<br>DDPKAVRATTPFKELGFDSLAAVRLRNLLNAATGLRLPSTLVFDHPNASAVAGFLTSELGSGTPAREASSALRDGYRQAGV<br>SGRVRSYLDLLAGLSDFREHFDGSDGFSLDLVDMADGPGEVTVICCAGTAAISGPHEFTRLAGALRGIAPVRAVPQPGYEEG<br>EPLPSSMAAVAAVQADAVIRTQGDKPFVVAGHSAGALMAYALATELLDRGHPPRGVVLIDVYPPGHQDAMNAWLEELTA |

| | |
|---|---|
| | TLFDRETVRMDDTRLTALGAYDRLTGQWRPRETGLPTLLVSAGEPMGPWPDDSWKPTWPFEHDTVAVPGDHFTMVQEHA<br>DAIARHIDAWLGGGNSKLAAALEHHHHHH* |
| MK149<br>SZ4-L-M3-TE | MQKVAELKNRVAVKLNRNEQLKNKVEELKNRNAYLKNELATLENEVARLENDVAEGGSGGGSGKLDPIAIVSMACRLPG<br>GVNTPQRLWELLREGGETLSGFPTDRGWDLARLHHPDPDNPGTSYVDKGGFLDDAAGFDAEFFGVSPREAAAMDPQQRLL<br>LETSWELVENAGIDPHSLRGTATGVFLGVAKFGYGEDTAAAEDVEGYSVTGVAPAVASGRISYTMGLEGPSISVDTACSSSL<br>VALHLAVESLRKGESSMAVVGGAAVMATPGVFVDFSRQRALAADGRSKAFGAGADGFGFSEGVTLVLLERLSEARRNGH<br>EVLAVVRGSALNQDGASNGLSAPSGPAQRRVIRQALESCGLEPGDVDAVEAHGTGTALGDPIEANALLDTYGRDRDADRP<br>LWLGSVKSNIGHTQAAAGVTGLLKVVLALRNGELPATLHVEEPTPHVDWSSGGVALLAGNQPWRRGERTRRARVSAFGIS<br>GTNAHVIVEEAPEREHRETTAHDGRPVPLVVSARTTAALRAQAAQIAELLERPDADLAGVGLGLATTRARHEHRAAVVAST<br>REEAVRGLREIAAGAATADAVVEGVTEVDGRNVVFLFPGQGSQWAGMGAELLSSSPVFAGKIRACDESMAPMQDWKVSD<br>VLRQAPGAPGLDRVDVVQPVLFAVMVSLAELWRSYGVEPAAVVGHSQGEIAAAHVAGALTLEDAAKLVVGRSRLMRSLS<br>GEGGMAAVALGEAAVRERLRPWQDRLSVAAVNGPRSVVVSGEPGALRAFSEDCAAEGIRVRDIDVDYASHSPQIERVREE<br>LLETTGDIAPRPARVTFHSTVESRSMDGTELDARYWYRNLRETVRFADAVTRLAESGYDAFIEVSPHPVVVQAVEEAVEEA<br>DGAEDAVVVGSLHRDGGDLSAFLRSMATAHVSGVDIRWDVALPGAAPFALPTYPFQRKRYWLQPAAPAAASDELAYRVS<br>WTPIEKPESGNLDGDWLVVTPLISPEWTEMLCEAINANGGRALRCEVDTSASRTEMAQAVAQAGTGFRGVLSLLSSDESAC<br>RPGVPAGAVGLLTLVQALGDAGVDAPVWCLTQGAVRTPADDDLARPAQTTAHGFAQVAGLELPGRWGGVVDLPESVDD<br>AALRLLVAVLRGGGRAEDHLAVRDGRLHGRRVVRASLPQSGSRSWTPHGTVLVTGAASPVGDQLVRWLADRGAERLVLA<br>GACPGDDLLAAVEEAGASAVVCAQDAAALREALGDEPVTALVHAGTLTNFGSISEVAPEEFAETIAAKTALLAVLDEVLGD<br>RAVEREVYCSSVAGIWGGAGMAAYAAGSAYLDALAEHHRARGRSCTSVAWTPWALPGGAVDDGYLRERGLRSLSADRA<br>MRTWERVLAAGPVSVAVADVDWPVLSEGFAATRPTALFAELAGRGGQAEAEPDSGPTGEPAQRLAGLSPDEQQENLLELV<br>ANAVAEVLGHESAAEINVRRAFSELGLDSLNAMALRKRLSASTGLRLPASLVFDHPTVTALAQHLTSQLDSGTPAREASSAL<br>RDGYRQAGVSGRVRSYLDLLAGLSDFREHFDGSDGFSLDLVDMADGPGEVTVICCAGTAAISGPHEFTRLAGALRGIAPVR<br>AVPQPGYEEGEPLPSSMAAVAAVQADAVIRTQGDKPFVVAGHSAGALMAYALATELLDRGHPPRGVVLIDVYPPGHQDA<br>MNAWLEELTATLFDRETVRMDDTRLTALGAYDRLTGQWRPRETGLPTLLVSAGEPMGPWPDDSWKPTWPFEHDTVAVPG<br>DHFTMVQEHADAIARHIDAWLGGGNSKLAAALEHHHHHH* |
| MK150<br>(5)M1-L-SZ3 | MSGDNGMTEEKLRRYLKRTVTELDSVTARLREVEHRAGEPVAVVAMACRLPGGVSTPEEFWELLSEGRDAVAGLPTDRG<br>WDLDSLFHPDPTRSGTAHQRGGGFLTEATAFDPAFFGMSPREALAVDPQQRLMLELSWEVLERAGIPPTSLQASPTGVFVGL<br>IPQEYGPRLAEGGEGVEGYLMTGTTTSVASGRIAYTLGLEGPAISVDTACSSSLVAVHLACQSLRRGESSLAMAGGVTVMPT<br>PGMLVDFSRMNSLAPDGRCKAFSAGANGFGMAEGAGMLLLERLSDARRNGHPVLAVLRGTAVNSDGASNGLSAPNGRAQ<br>VRVIQQALAESGLGPADIDAVEAHGTGTRLGDPIEARALFEAYGRDREQPLHLGSVKSNLGHTQAAAGVAGVIKMVLAMR<br>AGTLPRTLHASERSKEIDWSSGAISLLDEPEPWPAGARPRRAGVSSFGISGTNAHAIIEEAPQVVEGERVEAGDVVAPWVLSA<br>SSAEGLRAQAARLAAHLREHPGQDPRDIAYSLATGRAALPHRAAFAPVDESAALRVLDGLATGNADGAAVGTSRAQQRAV<br>FVFPGQGWQWAGMAVDLLDTSPVFAAALRECADALEPHLDFEVIPFLRAEAARREQDAALSTERVDVVQPVMFAVMVSL<br>ASMWRAHGVEPAAVIGHSQGEIAAACVAGALSLDDAARVVALRSRVIATMPGNKGMASIAAPAGEVRARIGDRVEIAAVN<br>GPRSVVVAGDSDELDRLVASCTTECIRAKRLAVDYASHSSHVETIRDALHAELGEDFHPLPGFVPFFSTVTGRWTQPDELDA<br>GYWYRNLRRTVRFADAVRALAEQGYRTFLEVSAHPILTAAIEEIGDGSGADLSAIHSLRRGDGSLADFGEALSRAFAAGVA<br>VDWESVHLGTGARRVPLPTYPFQRERVWLEPKPVARRSTEVDEVSALRYRIEWRPTGAGEPARLDGTWLVAKYAGTADET<br>STAAREALESAGARVRELVVDARCGRDELAERLRSVGEVAGVLSLLAVDEAEPEEAPLALASLADTLSLVQAMVSAELGCP<br>LWTVTESAVATGPFERVRNAAHGALWGVGRVIALENPAVWGGLVDVPAGSVAELARHLAAVVSGGAGEDQLALRADGV<br>YGRRWVRAAAPATDDEWKPTGTVLVTGGTGGVGGQIARWLARRGAPHLLLVSRSGPDADGAGELVAELEALGARTTVA |

| | |
|---|---|
| | ACDVTDRESVRELLGGIGDDVPLSAVFHAAATLDDGTVDTLTGERIERASRAKVLGARNLHELTRELDLTAFVLFSSFASAF<br>GAPGLGGYAPGNAYLDGLAQQRRSDGLPATAVAWGTWAGSGMAEGPVADRFRRHGVIEMPPETACRALQNALDRAEVC<br>PIVIDVRWDRFLLAYTAQRPTRLFDEIDDARRAAPQAAAEPRVGALASLPAPEREKALFELVRSHAAAVLGHASAERVPAD<br>QAFAELGVDSLSALELRNRLGAATGVRLPTTTVFDHPDVRTLAAHLAAELGGGSGGGSGNEVTTLENDAAFIENENAYLEK<br>EIARLRKEKAALRNRLAHKKLEHHHHHH* |
| MK152<br>SZ4-L-KR1-ACP1-<br>$L_{12}$-M3-TE | MQKVAELKNRVAVKLNRNEQLKNKVEELKNRNAYLKNELATLENEVARLENDVAEGGSGGGSGDEVSALRYRIEWRPTG<br>AGEPARLDGTWLVAKYAGTADETSTAAREALESAGARVRELVVDARCGRDELAERLRSVGEVAGVLSLLAVDEAEPEEAP<br>LALASLADTLSLVQAMVSAELGCPLWTVTESAVATGPFERVRNAAHGALWGVGRVIALENPAVWGGLVDVPAGSVAELA<br>RHLAAVVSGGAGEDQLALRADGVYGRRWVRAAAPATDDEWKPTGTVLVTGGTGGVGGQIARWLARRGAPHLLLVSRSG<br>PDADGAGELVAELEALGARTTVAACDVTDRESVRELLGGIGDDVPLSAVFHAAATLDDGTVDTLTGERIERASRAKVLGA<br>RNLHELTRELDLTAFVLFSSFASAFGAPGLGGYAPGNAYLDGLAQQRRSDGLPATAVAWGTWAGSGMAEGPVADRFRRH<br>GVIEMPPETACRALQNALDRAEVCPIVIDVRWDRFLLAYTAQRPTRLFDEIDDARRAAPQAAAEPRVGAHMLASLPAPERE<br>KALFELVRSHAAAVLGHASAERVPADQAFAELGVDSLSALELRNRLGAATGVRLPTTTVFDHPDVRTLAAHLAAELGGAT<br>GAEQAAPATTAPVDDPIAIVSMACRLPGGVNTPQRLWELLREGGETLSGFPTDRGWDLARLHHPDPDNPGTSYVDKGGFLD<br>DAAGFDAEFFGVSPREAAAMDPQQRLLLETSWELVENAGIDPHSLRGTATGVFLGVAKFGYGEDTAAAEDVEGYSVTGVA<br>PAVASGRISYTMGLEGPSISVDTACSSSLVALHLAVESLRKGESSMAVVGGAAVMATPGVFVDFSRQRALAADGRSKAFGA<br>GADGFGFSEGVTLVLLERLSEARRNGHEVLAVVRGSALNQDGASNGLSAPSGPAQRRVIRQALESCGLEPGDVDAVEAHGT<br>GTALGDPIEANALLDTYGRDRDADRPLWLGSVKSNIGHTQAAAGVTGLLKVVLALRNGELPATLHVEEPTPHVDWSSGGV<br>ALLAGNQPWRRGERTRRARVSAFGISGTNAHVIVEEAPEREHRETTAHDGRPVPLVVSARTTAALRAQAAQIAELLERPDA<br>DLAGVGLGLATTRARHEHRAAVVASTREEAVRGLREIAAGAATADAVVEGVTEVDGRNVVFLFPGQGSQWAGMGAELLS<br>SSPVFAGKIRACDESMAPMQDWKVSDVLRQAPGAPGLDRVDVVQPVLFAVMVSLAELWRSYGVEPAAVVGHSQGEIAAA<br>HVAGALTLEDAAKLVVGRSRLMRSLSGEGGMAAVALGEAAVRERLRPWQDRLSVAAVNGPRSVVVSGEPGALRAFSEDC<br>AAEGIRVRDIDVDYASHSPQIERVREELLETTGDIAPRPARVTFHSTVESRSMDGTELDARYWYRNLRETVRFADAVTRLAE<br>SGYDAFIEVSPHPVVVQAVEEAVEEADGAEDAVVVGSLHRDGGDLSAFLRSMATAHVSGVDIRWDVALPGAAPFALPTYP<br>FQRKRYWLQPAAPAAASDELAYRVSWTPIEKPESGNLDGDWLVVTPLISPEWTEMLCEAINANGGRALRCEVDTSASRTE<br>MAQAVAQAGTGFRGVLSLLSSDESACRPGVPAGAVGLLTLVQALGDAGVDAPVWCLTQGAVRTPADDDLARPAQTTAHG<br>FAQVAGLELPGRWGGVVDLPESVDDAALRLLVAVLRGGGRAEDHLAVRDGRLHGRRVVRASLPQSGSRSWTPHGTVLVT<br>GAASPVGDQLVRWLADRGAERLVLAGACPGDDLLAAVEEAGASAVVCAQDAAALREALGDEPVTALVHAGTLTNFGSIS<br>EVAPEEFAETIAAKTALLAVLDEVLGDRAVEREVYCSSVAGIWGGAGMAAYAAGSAYLDALAEHHRARGRSCTSVAWTP<br>WALPGGAVDDGYLRERGLRSLSADRAMRTWERVLAAGPVSVAVADVDWPVLSEGFAATRPTALFAELAGRGGQAEAEP<br>DSGPTGEPAQRLAGLSPDEQQENLLELVANAVAEVLGHESAAEINVRRAFSELGLDSLNAMALRKRLSASTGLRLPASLVFD<br>HPTVTALAQHLTSQLDSGTPAREASSALRDGYRQAGVSGRVRSYLDLLAGLSDFREHFDGSDGFSLDLVDMADGPGEVTVI<br>CCAGTAAISGPHEFTRLAGALRGIAPVRAVPQPGYEEGEPLPSSMAAVAAVQADAVIRTQGDKPFVVAGHSAGALMAYAL<br>ATELLDRGHPPRGVVLIDVYPPGHQDAMNAWLEELTATLFDRETVRMDDTRLTALGAYDRLTGQWRPRETGLPTLLVSAG<br>EPMGPWPDDSWKPTWPFEHDTVAVPGDHFTMVQEHADAIARHIDAWLGGGNSSSVDKLAAALEHHHHHH* |
| MK168<br>SZ4-L-KR5-ACP5(2) | MQKVAELKNRVAVKLNRNEQLKNKVEELKNRNAYLKNELATLENEVARLENDVAEGGSGGGSGPIPTGGRARDEDDDW<br>RYQVVWREAEWESASLAGRVLLVTGPGVPSELSDAIRSGLEQSGATVLTCDVESRSTIGTALEAADTDALSTVVSLLSRDGE<br>AVDPSLDALALVQALGAAGVEAPLWVLTRNAVQVADGELVDPAQAMVGGLGRVVGIEQPGRWGGLVDLVDADAASIRS<br>LAAVLADPRGEEQVAIRADGIKVARLVPAPARAARTRWSPRGTVLVTGGTGGIGAHVARWLARSGAEHLVLLGRRGADAP<br>GASELREELTALGTGVTIAACDVADRARLEAVLAAERAEGRTVSAVMHAAGVSTSTPLDDLTEAEFTEIADVKVRGTVNLD |

| | |
|---|---|
| | ELCPDLDAFVLFSSNAGVWGSPGLASYAAANAFLDGFARRRRSEGAPVTSIAWGLWAGQNMAGDEGGEYLRSQGLRAMD<br>PDRAVEELHITLDHGQTSVSVVDMDRRRFVELFTAARHRPLFDEIAGARAEARQSEEGPALAQRLAALSTAERREHLAHLIR<br>AEVAAVLGHGDDAAIDRDRAFRDLGFDSMTAVDLRNRLAAVTGVREAATVVFDHPTITRLADHYLERLVGTEVRGEAPSA<br>LAGLDALEAALPEVPATEREELVQRLERMLAALRPVAQAADASGTGANPSGDDLGEAGVDELLEALGRELDGDPNSSSVD<br>KLAAALEHHHHHH* |
| MK169<br>SZ4-L-RIFS(KR1-ACP)(2) | MQKVAELKNRVAVKLNRNEQLKNKVEELKNRNAYLKNELATLENEVARLENDVAEGGSGGGSGEPAEPASAGDPLLGTV<br>VSTPGSDRLTAVAQWSRRAQPWAVDGLVPNAALVEAAIRLGDLAGTPVVGELVVDAPVVLPRRGSREVQLIVGEPGEQRR<br>RPIEVFSREADEPWTRHAHGTLAPAAAAVPEPAAAGDATDVTVAGLRDADRYGIHPALLDAAVRTVVGDDLLPSVWTGVS<br>LLASGATAVTVTPTATGLRLTDPAGQPVLTVESVRGTPFVAEQGTTDALFRVDWPEIPLPTAETADFLPYEATSAEATLSAL<br>QAWLADPAETRLAVVTGDCTEPGAAAIWGLVRSAQSEHPGRIVLADLDDPAVLPAVVASGEPQVRVRNGVASVPRLTRVT<br>PRQDARPLDPEGTVLITGGTGTLGALTARHLVTAHGVRHLVLVSRRGEAPELQEELTALGASVAIAACDVADRAQLEAVLR<br>AIPAEHPLTAVIHTAGVLDDGVVTELTPDRLATVRRPKVDAARLLDELTREADLAAFVLFSSAAGVLGNPGQAGYAAANAE<br>LDALARQRNSLDLPAVSIAWGYWATVSGMTEHLGDADLRRNQRIGMSGLPADEGMALLDAAIATGGTLVAAKFDVAALR<br>ATAKAGGPVPPLLRGLAPLPRRAAAKTASLTERLAGLAETEQAAALLDLVRRHAAEVLGHSGAESVHSGRTFKDAGFDSLT<br>AVELRNRLAAATGLTLSPAMIFDYPKPPALADHLRAKLFGTEVRGEAPSALAGLDALEAALPEVPATEREELVQRLERMLA<br>ALRPVAQAADASGTGANPSGDDLGEAGVDELLEALGRELDGDPNSSSVDKLAAALEHHHHHH* |
| MK170<br>SZ4-L-M5-TE | MQKVAELKNRVAVKLNRNEQLKNKVEELKNRNAYLKNELATLENEVARLENDVAEGGSGGGSGKLEPIAIVGMACRFPG<br>DVDSPESFWEFVSGGGDAIAEAPADRGWEPDPDARLGGMLAAAGDFDAGFFGISPREALAMDPQQRIMLEISWEALERAG<br>HDPVSLRGSATGVFTGVGTVDYGPRPDEAPDEVLGYVGTGTASSVASGRVAYCLGLEGPAMTVDTACSSGLTALHLAMES<br>LRRDECGLALAGGVTVMSSPGAFTEFRSQGGLAADGRCKPFSKAADGFGLAEGAGVLVLQRLSAARREGRPVLAVLRGSA<br>VNQDGASNGLTAPSGPAQQRVIRRALENAGVRAGDVDYVEAHGTGTRLGDPIEVHALLSTYGAERDPDDPLWIGSVKSNIG<br>HTQAAAGVAGVMKAVLALRHGEMPRTLHFDEPSPQIEWDLGAVSVVSQARSWPAGERPRRAGVSSFGISGTNAHVIVEEA<br>PEADEPEPAPDSGPVPLVLSGRDEQAMRAQAGRLADHLAREPRNSLRDTGFTLATRRSAWEHRAVVVGDRDDALAGLRAV<br>ADGRIADRTATGQARTRRGVAMVFPGQGAQWQGMARDLLRESQVFADSIRDCERALAPHVDWSLTDLLSGARPLDRVDV<br>VQPALFAVMVSLAALWRSHGVEPAAVVGHSQGEIAAAHVAGALTLEDAAKLVAVRSRVLRRLGGQGGMASFGLGTEQA<br>AERIGRFAGALSIASVNGPRSVVVAGESGPLDELIAECEAEGITARRIPVDYASHSPQVESLREELLTELAGISPVSADVALYST<br>TTGQPIDTATMDTAYWYANLREQVRFQDATRQLAEAGFDAFVEVSPHPVLTVGIEATLDSALPADAGACVVGTLRRDRGG<br>LADFHTALGEAYAQGVEVDWSPAFADARPVELPVYPFQRQRYWLPIPTGGRARDEDDDWRYQVVWREAEWESASLAGR<br>VLLVTGPGVPSELSDAIRSGLEQSGATVLTCDVESRSTIGTALEAADTDALSTVVSLLSRDGEAVDPSLDALALVQALGAAG<br>VEAPLWVLTRNAVQVADGELVDPAQAMVGGLGRVVGIEQPGRWGGLVDLVDADAASIRSLAAVLADPRGEEQVAIRADG<br>IKVARLVPAPARAARTRWSPRGTVLVTGGTGGIGAHVARWLARSGAEHLVLLGRRGADAPGASELREELTALGTGVTIAA<br>CDVADRARLEAVLAAERAEGRTVSAVMHAAGVSTSTPLDDLTEAEFTEIADVKVRGTVNLDELCPDLDAFVLFSSNAGVW<br>GSPGLASYAAANAFLDGFARRRRSEGAPVTSIAWGLWAGQNMAGDEGGEYLRSQGLRAMDPDRAVEELHITLDHGQTSV<br>SVVDMDRRRFVELFTAARHRPLFDEIAGARAEARQSEEGPALAQRLAALSTAERREHLAHLIRAEVAAVLGHGDDAAIDRD<br>RAFRDLGFDSMTAVDLRNRLAAVTGVREAATVVFDHPTITRLADHYLERLVSGTPAREASSALRDGYRQAGVSGRVRSYL<br>DLLAGLSDFREHFDGSDGFSLDLVDMADGPGEVTVICCAGTAAISGPHEFTRLAGALRGIAPVRAVPQPGYEEGEPLPSSMA<br>AVAAVQADAVIRTQGDKPFVVAGHSAGALMAYALATELLDRGHPPRGVVLIDVYPPGHQDAMNAWLEELTATLFDRETV<br>RMDDTRLTALGAYDRLTGQWRPRETGLPTLLVSAGEPMGPWPDDSWKPTWPFEHDTVAVPGDHFTMVQEHADAIARHID<br>AWLGGGNSKLAAALEHHHHHH* |

**Table S3: Yields of proteins used in this study.** If no indication is given the PKS domain/modules are derived from DEBS. Typical yields are presented.

| Construct | Protein | Yield /mg/L of culture |
|---|---|---|
| MK96 | KR1-ACP1(2) | 8.6 |
| MK111 | (5)KS1-AT1-SZ3 | 1.3 |
| MK112 | SZ4-KR1-ACP1(2) | 7.9 |
| MK113 | (5)KS1-AT1 | 1.1 |
| MK136 | SZ4-KR1-ACP1-M2-TE | 0.9 |
| MK137 | SZ4-KR5-ACP5-M6-TE | 2.1 |
| MK138 | SZ4-KR1-ACP1-M2-TE | 3.4 |
| MK141 | SZ4-KR1-ACP1-M5-TE | 1.6 |
| MK142 | SZ4-KR1-ACP1-M6-TE | 1.4 |
| MK147 | SZ4-M6-TE | 7.9 |
| MK148 | SZ4-M2-TE | 1.5 |
| MK149 | SZ4-M3-TE | 3.8 |
| MK150 | (5)M1-SZ3 | 15.3 |
| MK152 | SZ4-KR1-ACP1-M3-TE | 3.6 |
| MK168 | SZ4-KR5-ACP5(2) | 2.3 |
| MK169 | SZ4-RIFS(KR1-ACP1)(2) | 9.8 |
| MK170 | SZ4-M5-TE | 3.3 |
| BL12 | LDD(4) | 6.7 |
| BL13 | (5)M1(2) | 2.4 |
| BL16 | (3)M2-TE | 1.6 |
| BL17 | (3)M5-TE | 0.8 |
| BL18 | (3)M6-TE | 4.9 |
| RSG34 | (3)M3-TE | 9.4 |
